# STING dampens the unfolded protein response to enable the presentation of self-antigens on MHC-I during inflammation

**DOI:** 10.64898/2026.05.25.727656

**Authors:** Ahmed M. Fahmy, Ali Ahmadi, Joël Lanoix, Tyler Cannon, Moustafa N. Elemeery, Camberly Hernandez Paredes, Benoit Barrette, Eric Bonneil, Yong Zhong Xu, Maha Ibrahim, Guillermo Arango-Duque, Eric Olivier Audemard, Sébastien Lemieux, Eric Chevet, Erwin Schurr, Philippe Pierre, Samantha Gruenheid, Pierre Thibault, Heidi M. McBride, Michel Desjardins

## Abstract

A growing body of evidence supports the contribution of the long-lasting adaptive immune system in Parkinson’s disease (PD). We showed that the PD-associated protein PINK1 negatively regulates the presentation of mitochondrial antigens (MitAP) on MHC-I molecules. *In vivo* evidence indicated that MitAP activation in mice, in the absence of PINK1, led to cytotoxic CD8^+^ T cell stimulation and severe motor impairments, reversible by L-DOPA. We show here that following TLR4 activation, MitAP is engaged through a pathway involving cGAS-STING, which acts as a rheostat to dampen the unfolded protein response (UPR). Without STING, the stress response is amplified, leading to a translational attenuation that inhibits the expression of XBP1s, a transcription factor required for MitAP. STING activity also regulates the repertoire of peptides displayed at the cell surface during inflammation, highlighting a potential role in immunosurveillance. These findings establish STING and the UPR as key immune regulators targetable for therapeutic intervention during autoimmune diseases and PD.

## Introduction

Parkinson’s disease (PD) is a progressive disorder characterized by the loss of dopaminergic neurons (DNs) in the substantia nigra of the brain, leading to movement impairments ^1^. While the neurodegenerative nature of PD is well established, the molecular mechanisms responsible for the initiation of the disease, which begins several years before the onset of motor symptoms ^2,3^, are still elusive. This impairs the development of effective therapeutic approaches for the early stages of the disease, a key moment to intervene and limit the loss of DNs. Together with ageing, inflammation, a rapid innate immune response triggered during infection or tissue damage, has been identified as a key determinant for PD, pointing to a role of the immune system in the disease process ^4^. Furthermore, post-mortem analyses have shown that T lymphocytes (T cells) are present in the brain of people with PD and animal models of the disease ^5,6^, suggesting that adaptive immunity, the long-lasting arm of the immune system, also contributes to PD. In that context, the recent detection of autoreactive CD4^+^ T cells against a-synuclein peptides in people with PD emphasized the likelihood that autoimmune mechanisms play a role in PD pathophysiology ^7^.

The contribution of autoimmune mechanisms in the pathophysiological process leading to PD is supported by data obtained in cell systems and mouse models of the disease. It was shown that two PD-related proteins, PINK1 and Parkin, act as negative regulators in the presentation of antigens of mitochondrial origin (MitAP) on MHC class I molecules at the surface of antigen-presenting cells (APCs) during inflammation ^8^. Furthermore, gut infection in *Pink1^-/-^* mice was shown to engage MitAP and activate a population of autoreactive cytotoxic CD8^+^ T cells concomitant with the emergence of motor impairments fully reversible by L-DOPA treatment ^9^. Interestingly, a recent study reported that the depletion of CD8+ T cells in *Pink1^-/-^*mice before infection inhibited the motor impairments, linking T cells to the PD-like disease process ^10^. In a complementary study, the adoptive transfer in mice of cytotoxic CD8+ T cells specific for a mitochondrial antigen presented by the MitAP pathway was shown to induce motor symptoms with DNs destruction, providing further evidence for a direct role for MitAP and CD8+ T cells in the PD-like disease ^11^. While these studies underscored the significance of MitAP in PD pathogenesis, the cellular mechanisms leading to the presentation of mitochondrial antigens on MHC-I and the activation of cytotoxic CD8^+^ T cells, remain largely unknown.

A link between MitAP and PD in humans is emerging. The MitAP pathway is responsible for processing and presenting a peptide from the mitochondrial matrix protein 2-oxoglutarate dehydrogenase (OGDH). In humans, autoimmune mechanisms targeting OGDH peptides cause the disease Primary Biliary Cholangitis (PBC), also referred to as Primary Biliary Cirrhosis, which, like PD, is thought to arise from an interplay between underlying genetic predispositions and a second microbial trigger ^12^. Interestingly, PBC is effectively treated with Ursodeoxycholic acid (UDCA), a therapy recently shown to improve gait parameters in a small cohort of PD patients ^13^.

Here, using pharmacological and genetic approaches, we show that MitAP is a MyD88-independent process initiated by the TRAM/TRIF arm of TLR4 signaling. Remarkably, the stimulator of interferon genes (STING) plays a role in the activation of this antigen presentation pathway during inflammation by acting as a rheostat, controlling the magnitude of the unfolded protein response (UPR). STING’s involvement in coordinating the innate and adaptive immune responses shapes the repertoire of peptides (immunopeptidome) presented at the surface of macrophages and the activation of CD8+ T cells in inflammatory conditions. Characterizing the role played by the TLR4/STING/UPR axis in MitAP enhances our understanding of how autoimmune mechanisms may contribute to PD pathophysiology. It also identifies potential therapeutic targets along this axis to modulate the immune responses at the early stages of the disease, providing a new framework to explore the role of antigen presentation in neurodegenerative diseases and new avenues for translational research.

## Results

### TLR4 activates MitAP

We showed that MitAP is activated by Gram-negative bacteria and LPS, suggesting that TLR4 signaling modulates the engagement of this adaptive immune pathway during inflammation ^8^. To characterize the cellular pathways responsible for MitAP activation, we first compared the extent to which two Gram-negative bacteria, enteropathogenic *E. coli* (EPEC) and *Helicobacter pylori* (*H. pylori*), activated this pathway *in vitro* in wild-type (WT) RAW macrophages (Supp. Fig. 1A). To do so, we engineered a RAW macrophage cell line expressing the glycoprotein B (gB) of *Herpes virus* in the matrix of mitochondria ^8^. These cells were also modified to express the H-2K^b^ allele required to present a gB-derived peptide on MHC-I molecules at their surface. MitAP was monitored by measuring the activation (release of β-galactosidase) of a gB-specific CD8+ T cell hybridoma (2E2) co-incubated with fixed RAW cells (APCs). We observed that while EPEC effectively activated MitAP, *H. pylori* were unable to do so. Similar results were observed with heat-killed bacteria (Supp. Fig. 1B), ruling out the possibility that MitAP required an active microbial process, such as the secretion of microbial effectors in the cytoplasm. The lack of MitAP activation by *H. pylori* was likely because the structure of the LPS at the surface of this bacterium is unable to trigger TLR4 signaling ^14^. A confirmation that MitAP was initiated through TLR4 activation was provided by showing that incubation of cells with the TLR4 inhibitor TAK242 ^15^ before infection resulted in a strong dose-dependent inhibition of this pathway (Supp. Fig. 1C).

**Fig. 1:**
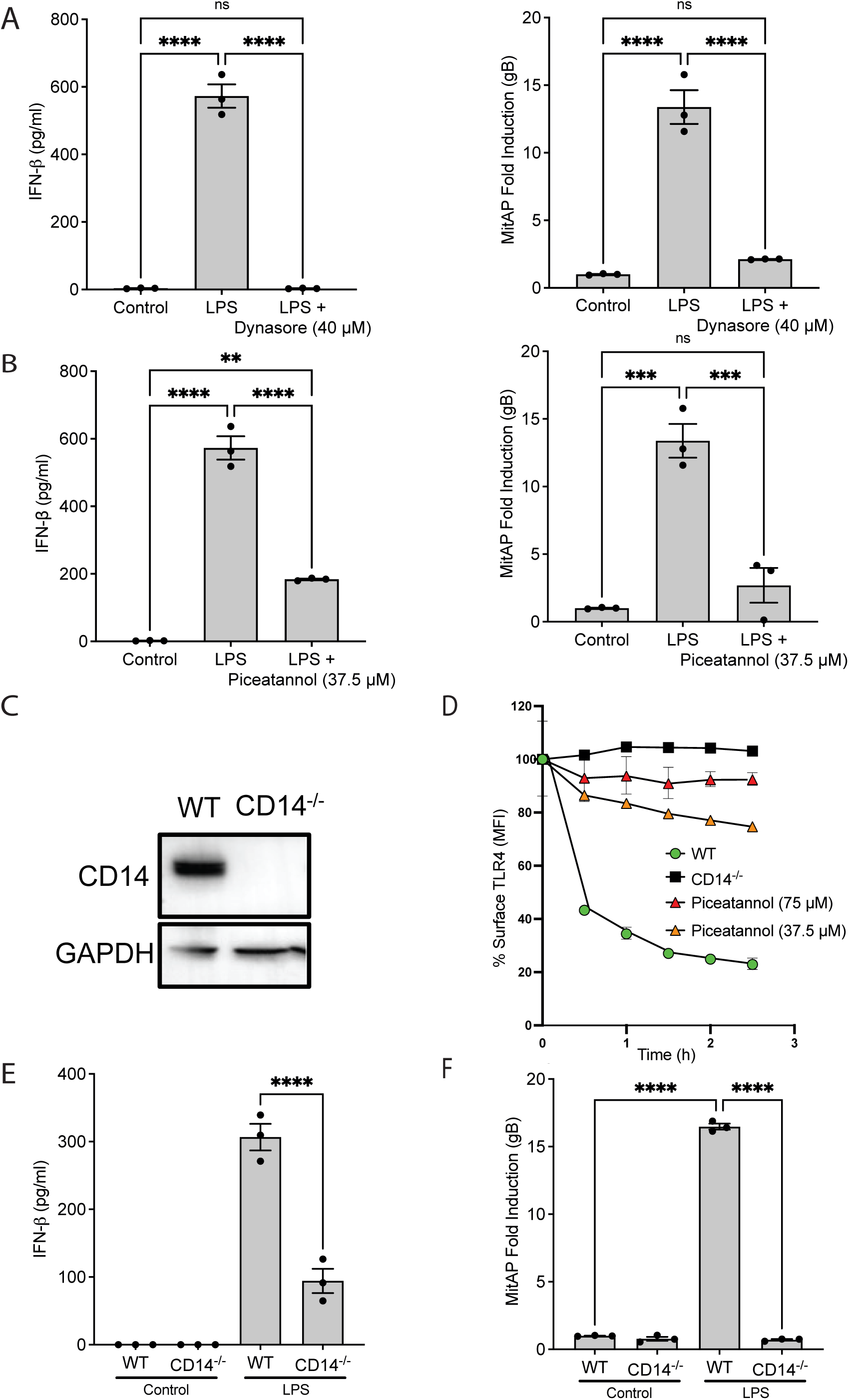
The TRAM/TRIF arm of TLR4 activates MitAP. **A**) Blocking endocytosis with Dynasore inhibits both IFN-β release and MitAP. **B**) Inhibition of CD14 internalization with the Syk inhibitor Piceatannol reduces both IFN-β release and MitAP. **C**) CD14 knockout (CD14^-/-^) RAW cells generated using CRISPR-Cas9 show loss of CD14 expression by Western blot analysis. **D**) Flow cytometry demonstrates impaired internalization of TLR4 from the cell surface upon LPS stimulation in CD14^-/-^ RAW cells. **E, F**) IFN-β release and MitAP are inhibited in CD14^-/-^ RAW cells.

We then measured the activation of MitAP by various microbial surface molecules. We observed that MitAP was activated in a concentration-dependent manner by LPS (Supp. Fig. 1D). In contrast, peptidoglycan and flagellin (recognized by TLR2 and TLR5, respectively) were poor inducers of MitAP; these molecules activating MitAP only at much higher concentrations (Supp. Fig. 1D). MitAP activation by LPS was completely inhibited when cells were pre-incubated with TAK242 (Supp. Fig. 1E), confirming the involvement of TLR4 in the presentation of mitochondrial antigens. On the other hand, TAK242 had no inhibitory effect when MitAP was induced with a high concentration of peptidoglycan and flagellin (Supp. Fig. 1F). These data suggest that the initial trigger of MitAP during infection is mediated through TLR signaling, with TLR4 playing a dominant role compared to TLR2 and TLR5.

### A non-canonical TRAM/TRIF arm-driven pathway activates MitAP

The binding of LPS to TLR4 engages two distinct signaling arms ^16^. The MyD88-dependent arm leads to the activation of NF-kB and the release of proinflammatory cytokines such as IL-6. Engagement of a second MyD88-independent arm involving the adaptor proteins TRAM and TRIF leads to the activation of IRF3 and the release of IFN-β. Treatment of RAW cells with a specific MyD88 inhibitory peptide before LPS stimulation effectively inhibited the release of IL-6 (Supp. Fig. 2A), demonstrating the efficacy of this inhibitory peptide. This peptide, however, had no inhibitory effect on the activation of MitAP by LPS (Supp. Fig. 2B). Similar results were obtained when the downstream effector NF-kB was inhibited with the specific inhibitor BAY11-7082 ^17^ (Supp. Fig. 2C and D). Accordingly, we concluded that the MyD88-dependent arm of TLR4 was not an essential player in MitAP activation.

**Fig. 2:**
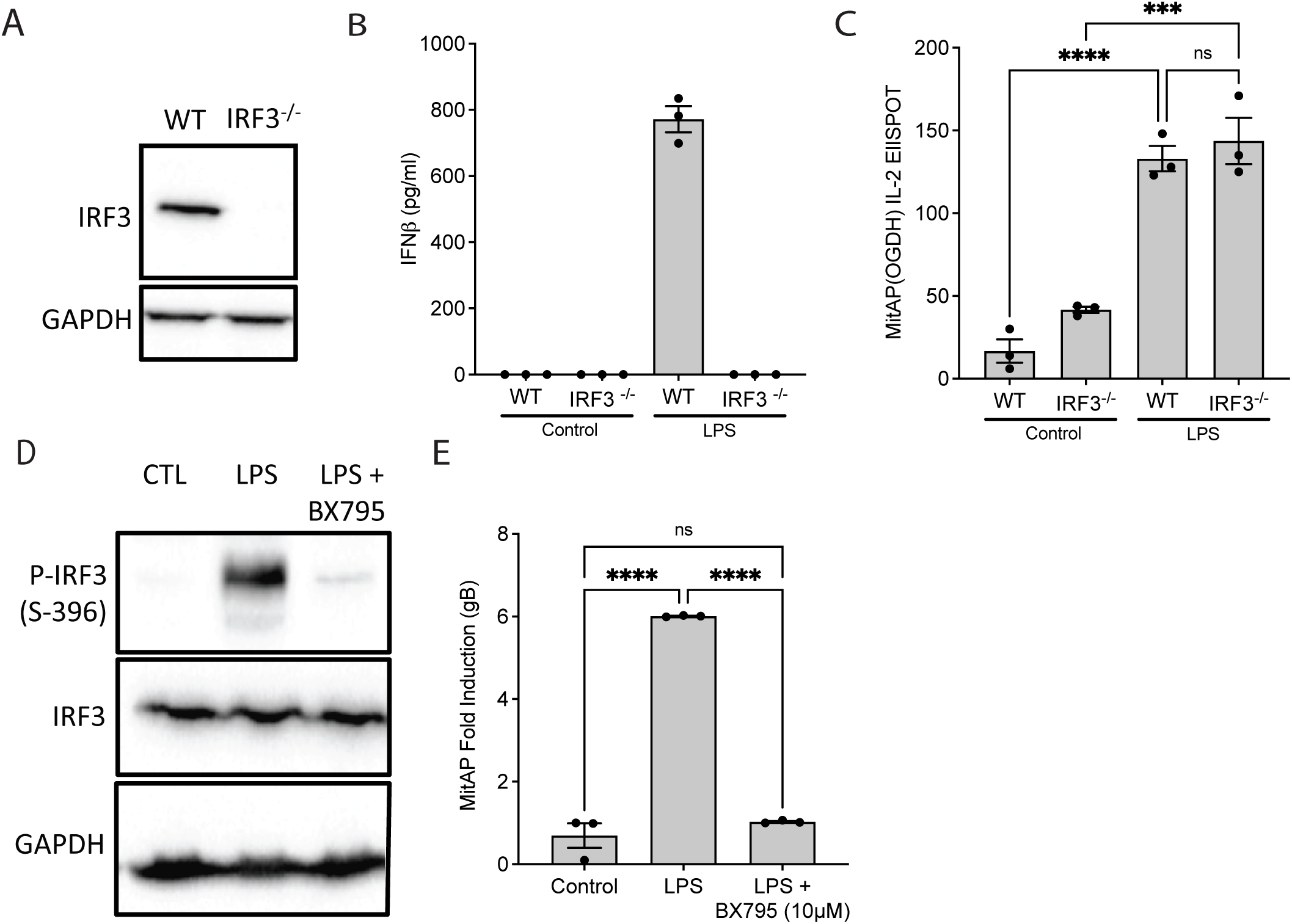
IRF3 is not required, but TBK1 is for MitAP activation. **A**) Loss of IRF3 expression in IRF3^-/-^ RAW cells generated via CRISPR-Cas9 was confirmed by Western blot analysis. **B)** The release of IFN-β in response to LPS is inhibited in IRF3^-/-^ RAW cells. **C**) MitAP activation by LPS is not affected by the loss of IRF3. **D**) Western blot analysis shows inhibition of phosphorylated IRF3 (P-IRF3), confirming the efficacy of the TBK1 inhibitor BX795. **E**) BX795-mediated inhibition of TBK1 suppresses MitAP activation.

The TRAM/TRIF arm is activated from endosomes following the internalization of TLR4 and its ligand by endocytosis ^18^. The finding that Dynasore, a molecule that inhibits endocytosis ^19^, strongly decreased the release of IFN-β (positive control) in response to LPS, as well as the induction of MitAP, suggested that the TRAM/TRIF arm of TLR4 was responsible for the activation of this antigen presentation pathway (Fig. 1A). Along the same line, pre-incubation of RAW cells with piceatannol, a molecule that targets more specifically the Syk kinase/CD14 complex required for TLR4 internalization^18^, also inhibited the release of IFN-β and MitAP in response to LPS (Fig. 1B). Finally, we generated CD14 knockout RAW cells (CD14^-/-^) using CRISPR-Cas9 (Fig. 1C) and observed that TLR4 was no longer internalized in response to LPS in these cells (Fig. 1D). Consequently, the release of IFN-β was inhibited (Fig. 1E) and MitAP was not induced (Fig. 1F). These data demonstrated that the TRAM/TRIF arm of TLR4 signaling was responsible for MitAP activation in inflammatory conditions.

The involvement of the TRAM/TRIF arm suggested that the transcription factor IRF3, a downstream effector along this arm of TLR4 signaling, was required for MitAP activation. To test this hypothesis, we generated an IRF3 KO RAW cell line (IRF3^-/-^) using CRISPR-Cas9 (Fig. 2A). Because the gB protein is not expressed in the IRF3^-/-^ cell line, MitAP was monitored by measuring the presentation of a peptide from an endogenous protein of the mitochondria matrix, OGDH, using a specific CD8+ T cell hybridoma (2CZ) in an ELISPOT assay. As anticipated, the release of IFN-β, whose expression is regulated by IRF3 ^20,21^, was decreased in IRF3^-/-^ cells in response to LPS (Fig. 2B). Interestingly, the induction of MitAP by LPS was not affected in IRF3^-/-^ cells (Fig. 2C), suggesting that a bifurcation within the TRAM/TRIF arm, upstream of IRF3, was responsible for MitAP activation.

A key molecular complex upstream of IRF3 is the kinase IKKe and its partner, the TANK-binding kinase-1 (TBK1), which phosphorylates IRF3 and facilitates its translocation to the nucleus ^22^. Having shown that IRF3 was not involved in MitAP, we tested whether the upstream protein TBK1 was involved. For this, we treated RAW cells with the specific TBK1 inhibitor BX795. As a positive control, we showed that incubation with BX795 before LPS treatment effectively blocked IRF3 phosphorylation (Fig. 2D). In these conditions, we observed that the treatment with BX795 resulted in a near-complete inhibition of MitAP (Fig. 2E). This indicates that following the activation of TLR4 by LPS, a non-canonical pathway involving the TRAM/TRIF arm and the TBK1/IKKe complex, distinct from IRF3 activation, was involved in MitAP.

### The cGAS-STING pathway is required for the activation of MitAP in inflammatory conditions

Given the critical importance of TBK1, but not the IRF3, we sought to understand how this protein could be involved in MitAP. In APCs, the production of type-I IFN molecules is regulated, at least in part, by the stimulator of interferon genes (STING), which interacts with TBK1 to activate IRF3 ^23^. The first indication that the cyclic GMP–AMP synthase (cGAS)–STING pathway was involved in the activation of MitAP came from showing that the addition of cGAMP to the culture medium, an agonist of STING, was as potent as LPS to induce this presentation pathway in RAW cells (Fig. 3A). cGAMP also activated MitAP in IRF3^-/-^ cells, indicating that the role of cGAS-STING in this presentation pathway occurred independently of IRF3 (Fig. 3B). We then used CRISPR-Cas9 to generate a STING KO in RAW cells (STING^-/-^) (Fig. 3C). We observed that MitAP was inhibited following LPS treatment in these cells, compared to WT cells (Fig. 3D).

**Fig. 3:**
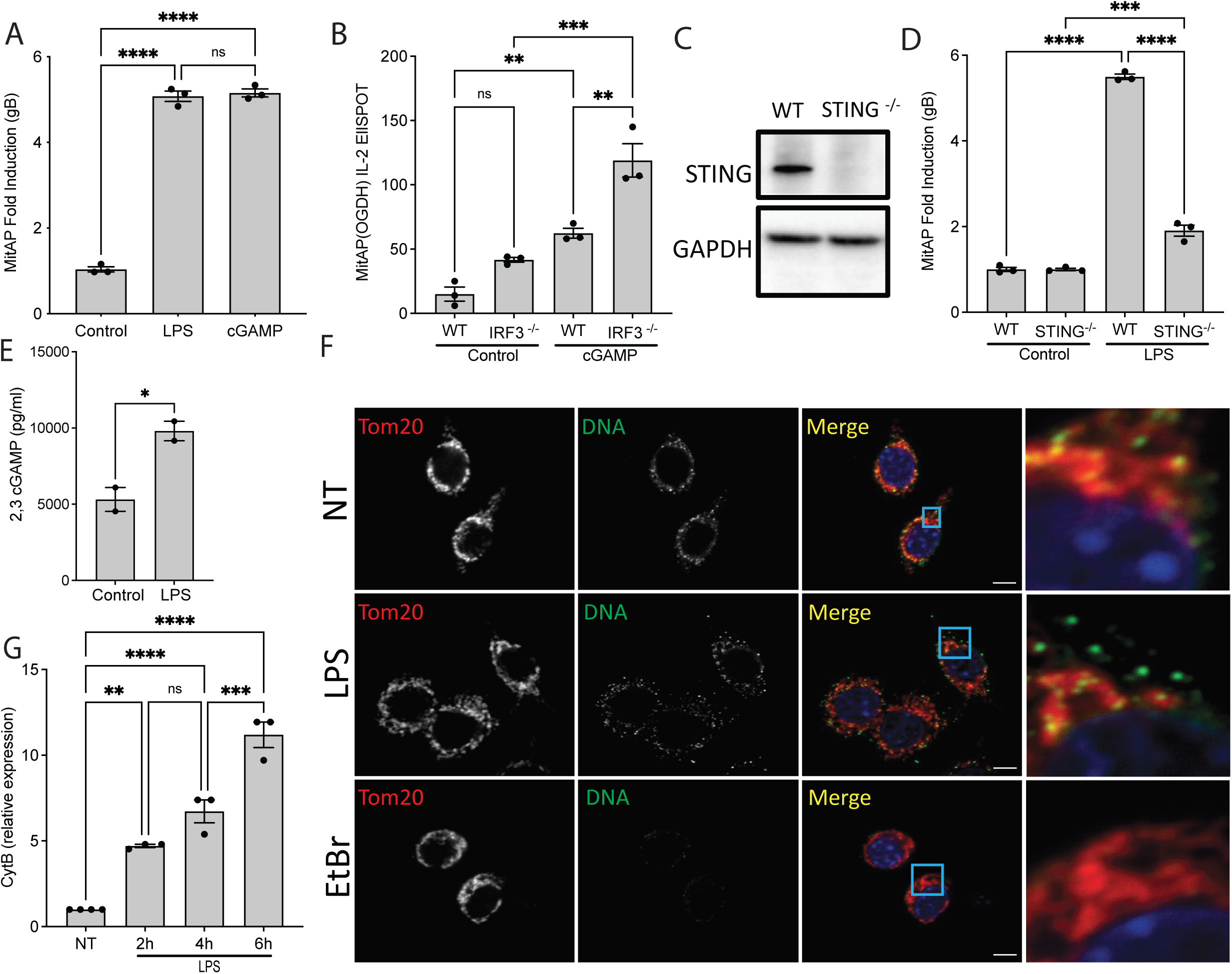
STING is required to activate MitAP along the TLR4-induced pathway. **A)** The STING agonist cGAMP induces MitAP. **B**) IRF3 is not required for cGAMP-mediated MitAP activation. **C**) Loss of STING expression in STING^-/-^ RAW cells generated via CRISPR-Cas9 was confirmed by Western blot analysis. **D**) MitAP activation in response to LPS is significantly inhibited in STING^-/-^ RAW cells. **E**) Treatment of RAW cells with LPS induces cGAMP production. **F**) Immunofluorescence microscopy shows that LPS treatment in RAW cells leads to the accumulation of mitochondrial DNA (mtDNA) within vesicular structures near mitochondria (TOM20). The DNA signal is absent after mtDNA depletion with ethidium bromide (EtBr) treatment (NT: non-treated). **G**) LPS treatment in RAW cells increases the level of mitochondrial DNA in the cytoplasm over time by qPCR using CytB-specific primers.

The cGAS-STING pathway is triggered by the presence of DNA in the cytosol, which binds to cGAS, enabling the production of cGAMP, which, in turn, binds to and activates STING in the endoplasmic reticulum (ER) ^24^. The observation that LPS treatment resulted in the production of cGAMP in WT RAW cells (Fig. 3E) suggested that DNA was released to the cytoplasm upon TLR4 activation. Although formally identifying the source of DNA activating cGAS-STING in response to LPS in our system was not the aim of our study, we observed that small vesicular-like structures containing DNA were present near mitochondria in LPS-treated cells (Fig. 3F). The specificity of the anti-DNA antibody was confirmed in ethidium bromide (EtBr)-treated RAW cells, where the selective depletion of mtDNA ^25^ resulted in the loss of the vesicular staining (Fig. 3F, lower panel). The vesicular structures revealed with the anti-DNA antibody are reminiscent of recently described mtDNA-containing mitochondria-derived vesicles (MDVs) involved in the transfer of mtDNA to the cytosol ^26^. Using qPCR, we measured increasing levels of mtDNA in the cytoplasm in response to LPS (Fig. 3G), suggesting that the DNA that activated cGAS-STING in our system was released from mitochondria, as observed in previous studies ^27,28^. While our data so far highlighted the sequential engagement of the TRAM/TRIF arm of TLR4 and the cGAS-STING pathway during MitAP activation by LPS, we still had to identify the molecular mechanisms at play downstream of STING, considering that its two main effectors, NFkB and IRF3 ^29^, were not essential for MitAP (Supp. Figs. 2D and 4B).

**Fig. 4:**
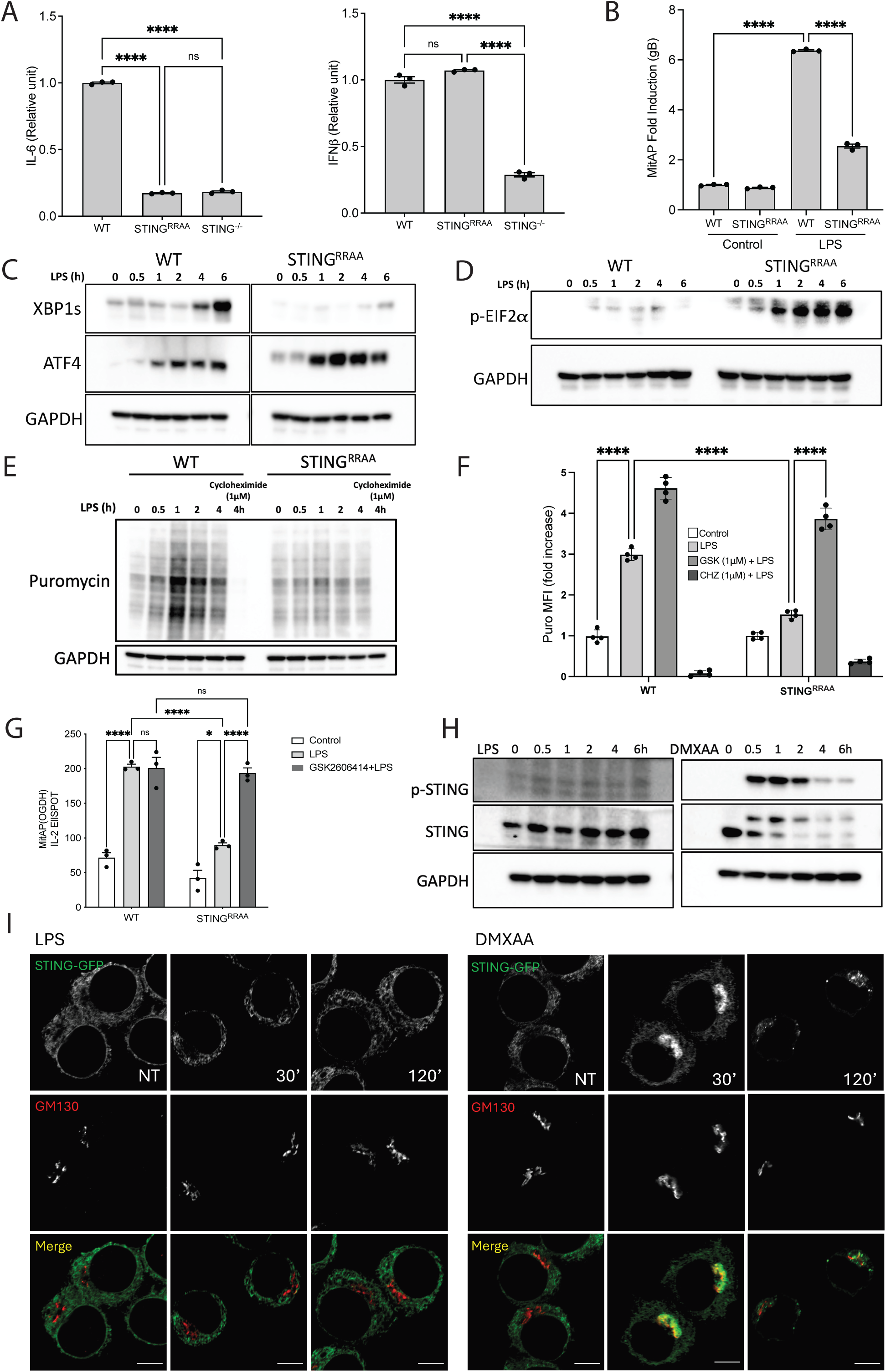
STING engages MitAP through UPR regulation. **A**) The secretion of IL-6 in response to LPS is inhibited in both STING^-/-^ and STING^RRAA^ cells, while the secretion of IFN-β is affected only in STING^-/-^ cells. **B**) MitAP is inhibited in STING^RRAA^ RAW cells in response to LPS. **C)** Both XBP1s and ATF4 are produced in response to LPS in WT RAW cells. In contrast, ATF4 is increased, and the production of XBP1s is strongly decreased in the STING^RRAA^ cells. **D**) Phosphorylation of eIF2a, a marker of UPR activation, is strongly increased in STING^RRAA^ cells. **E**) Western blotting with anti-puromycin antibodies indicates that protein synthesis is strongly decreased in the STING^RRAA^ cells. Protein synthesis is inhibited when cells are treated with Cycloheximide before LPS (negative control). **F**) The incorporation of puromycin in nascent polypeptides measured by FACS was recovered by treating the STING^RRAA^ cells with the PERK inhibitor before LPS treatment. **G**) MitAP was efficiently activated by LPS in STING^RRAA^ cells when PERK was inhibited with GSK2606414. **H**) Western blotting shows that in WT RAW cells STING is sequentially phosphorylated and degraded only in response to DMXAA treatment (p-STING: phosphorylated STING). **I**) In WT RAW cells expressing STING–GFP, LPS stimulation does not induce STING translocation to the Golgi, as assessed by anti-GFP staining and GM130 labelling. In contrast, treatment with the STING agonist DMXAA triggers a rapid translocation to the Golgi at 30’, followed by exit from the Golgi by 120’. Scale bar, 5 µm.

### Chemical inducers of the UPR activate MitAP

If not through NFkB and IRF3, how does STING regulate the activation of MitAP in inflammatory conditions? A recent study reported that a small cytoplasmic domain (referred to as the “UPR motif”) functionally connects STING to the unfolded protein response (UPR)^30^, suggesting that STING could regulate MitAP activation in coordination with the UPR. Accordingly, we first aimed to determine whether the engagement of the UPR activates MitAP. The UPR, a signaling mechanism triggered by the accumulation of improperly folded proteins in the lumen of the ER, has been shown to regulate cellular homeostasis and the inflammatory response ^31^. It is engaged by the activation of three sensors in the ER lumen, PERK, IRE1 and ATF6, leading to the production of ATF4, XBP1s and a cleaved form of ATF6. These proteins reach the nucleus and act as transcription factors regulating the expression of a wide array of proteins aimed at restoring ER homeostasis ^32^. The UPR can be induced in cells by chemicals that cause ER stress, including tunicamycin and thapsigargin (TG) ^33^. In RAW macrophages, treatment with both molecules led to robust activation of MitAP in a dose-dependent manner (Supp. Fig. 3A). We observed that the activation of MitAP by TG was not inhibited by the TLR4 inhibitor TAK242 (Supp. Fig. 3B) and still occurred in cells lacking CD14 (Supp. Fig. 3C), as well as when TBK1 was inhibited with BX795 (Supp. Fig. 3D). Thus, we concluded that TG activated MitAP downstream of the TLR4 signaling events occurring in the cytoplasm.

### STING acts in coordination with the UPR to engage MitAP in inflammatory conditions

We then conducted a series of experiments to determine whether a connection between STING and the UPR was at play for the activation of MitAP following TLR4 activation. For this, we took advantage of a recent study showing that a double R to A substitution in the “UPR motif” (at positions 331 and 334 in humans) affected the interaction between STING and the UPR in HEK293T cells, without affecting the activation of the IFN-β pathway and the expression of the interferon-stimulated gene (ISG) IFIT ^30^. Accordingly, we used CRISPR-Cas9 to edit the STING gene, introducing the two RRAA substitutions (position 330 and 333 in mice) in RAW cells (STING^RRAA^), arguing that this mutant would allow us to conclusively establish whether STING regulates MitAP through the IFN-β pathway (still active) or the UPR. As mentioned for the HEK293T cells above, we observed that in contrast with STING^-/-^ cells, where the secretion of both IL6 and IFN-β was inhibited, IFN-β was still secreted in the STING^RRAA^ macrophages (Fig. 4A). We then showed that MitAP was inhibited in STING^RRAA^ cells by measuring both the presentation of the gB (Fig. 4B) and OGDH peptides (Supp. Fig. 4). A second approach was used to monitor MitAP. Instead of detecting the presentation of the gB peptide with the hybridoma, we used a mass spectrometry-based approach to detect and measure the gB peptide loaded at the cell surface on MHC-I. To do so, the MHC-I-peptide complexes were immune-isolated with a mix of MHC-I-specific antibodies. Parallel Reaction Monitoring (PRM), a method to identify specific peptides in the mass spectrometer, confirmed that the presentation of the gB peptide associated with MitAP was upregulated in response to LPS in WT cells and downregulated in STING^-/-^ cells (Supp. Fig. 5). These data indicated that the IFN-β pathway was either not sufficient for or not involved in MitAP activation. They also showed that STING and the UPR interact along the MitAP pathway.

**Fig. 5:**
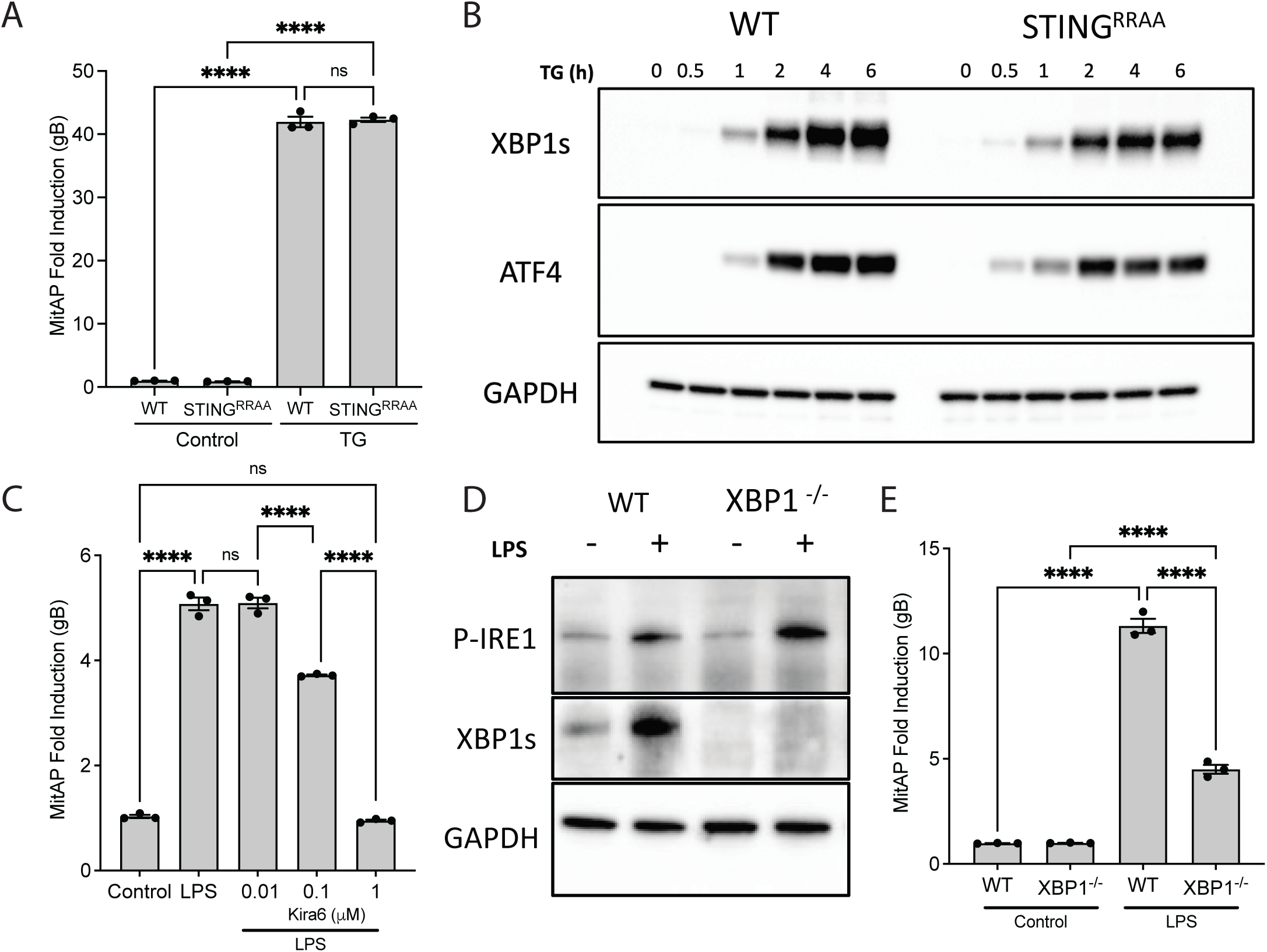
XBP1s is required for MitAP activation. **A**) TG treatment activates MitAP in STING^RRAA^ cells. **B**) Treatment with the UPR inducer TG induces the production of XBP1s and ATF4 in both WT and STING^RRAA^ cells. **C**) Inhibition of IRE1 with Kira6 before LPS treatment decreases MitAP in a dose-dependent manner. **D**) The expression of XBP1s in XBP1^-/-^ cells produced by CRISPR-Cas9 is inhibited. The phosphorylation of IRE1 (p-IRE1) by LPS, an event upstream of XBP1s production, is not affected in XBP1^-/-^ cells. **E**) MitAP activation in response to LPS is inhibited in XBP1^-/-^ cells.

Next, we sought to determine whether STING and the UPR act in coordination to activate MitAP. Hallmark steps in the UPR following the initial activation of PERK are the phosphorylation of eIF2a, which blocks cap-dependent translation and reduces protein synthesis ^34,35^. This block affects the expression of the transcription factor XBP1s but not ATF4. Thus, using Western blotting, we measured the expression of three essential sets of markers of the UPR after LPS treatment: 1) the expression of ATF4 and XBP1s, 2) the phosphorylation of eIF2a, and 3) the level of protein synthesis after puromycin labeling. In WT RAW cells, we observed that LPS treatment increased the levels of both XBP1s and ATF4 (Fig. 4C), indicating the ability of this molecule to activate the UPR, as shown previously ^36,37^. Kinetic analyses showed that ATF4 was detected as early as 30 min after LPS treatment, with an increase up to 4h post-treatment. XBP1s, on the other hand, was detected with a slightly delayed kinetics, as reported previously ^38^. Interestingly, LPS only induced a slight increase in the phosphorylation of eIF2a (Fig. 4D), without any apparent inhibition in protein synthesis, as assessed by puromycin incorporation into nascent polypeptides, which instead increased up to 2h after treatment (Fig. 4E), suggesting that an attenuated UPR was engaged. Next, we showed that the loss of the “UPR motif” in the STING^RRAA^ altered how cells activate the UPR in response to LPS. In these cells, we observed an increase in ATF4 production in response to LPS with a sharp decrease in the level of XBP1s expression (Fig. 4C). This was accompanied by a strong increase in the phosphorylation of eIF2a (Fig. 4D), and a concomitant decrease in protein synthesis (Fig. 4E), as described following UPR activation previously ^39^. The difference between XBP1s and ATF4 expression is probably because ATF4 mRNA translation is cap-independent ^40^. Thus, its expression is not affected by the eIF2a-induced translational attenuation. Inhibition of the translational attenuation, with the resulting increase in ATF4 and decrease in XBP1s expression, was also observed in the STING^-/-^cells (Supp. Fig. 6). Finally, further evidence that STING regulates the engagement of the UPR and MitAP upon DNA sensing was demonstrated by showing that both pathways were affected by the loss of cGAS in cGAS^-/-^ RAW cells (Supp. Fig. 7).

**Fig. 6:**
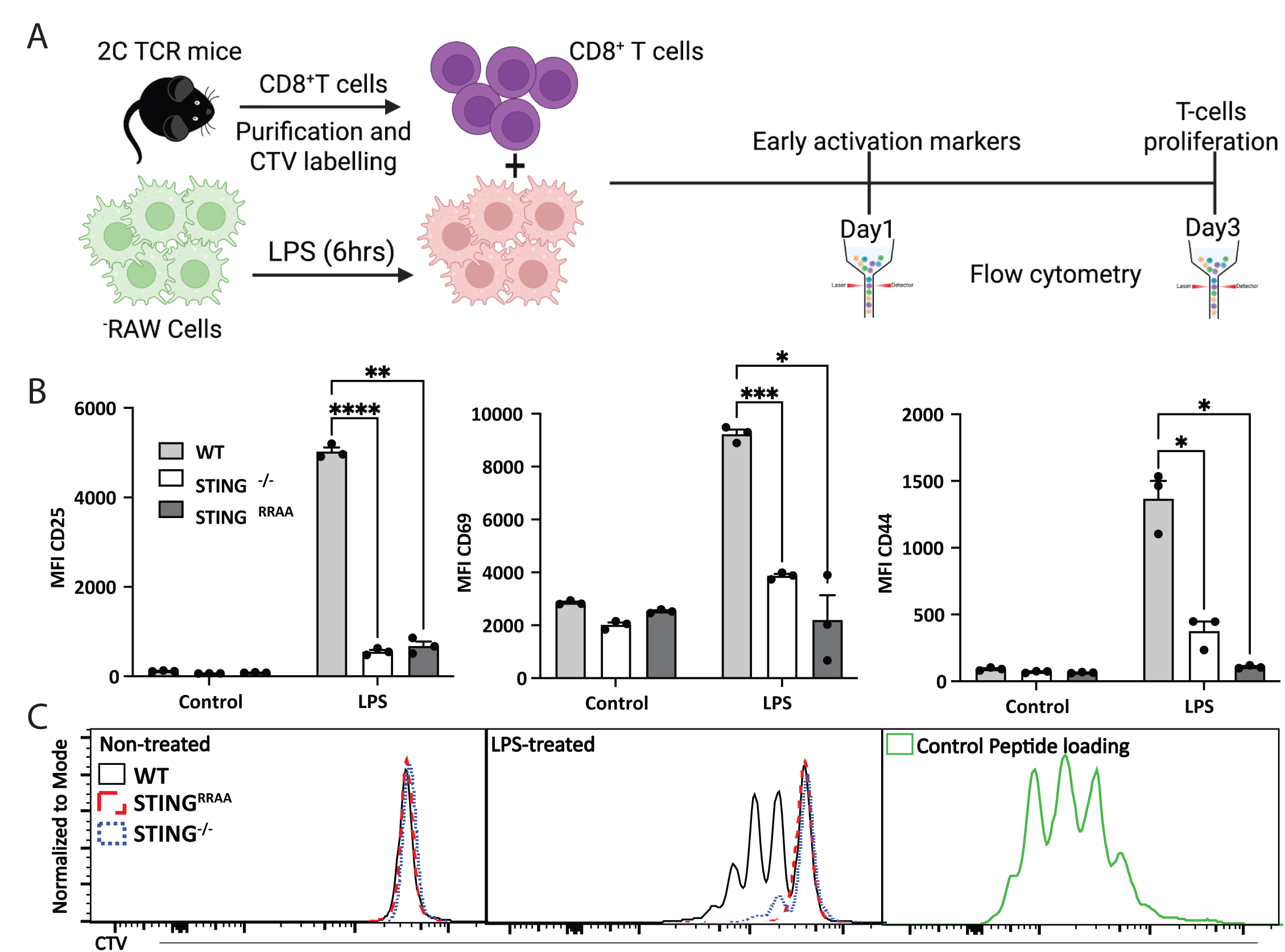
The STING pathway in APCs is required for 2C CD8^+^ T cell activation and proliferation. **A**) Schematic diagram illustrating the experimental workflow, including the methods and key time points. **B**) MFI quantification by flow cytometry demonstrates reduced activation of 2C CD8+ T cells, as indicated by the decrease of the activation markers CD25, CD44, and CD69 when the T cells are co-incubated with LPS-treated STING^-/-^ and STING^RRAA^ RAW cells. **C**) 2C CD8^+^T cell proliferation (assessed by flow cytometry using CTV staining) was suppressed when incubated with LPS-treated STING^-/-^ or STING^RRAA^ RAW cells, compared to WT cells.

**Fig. 7:**
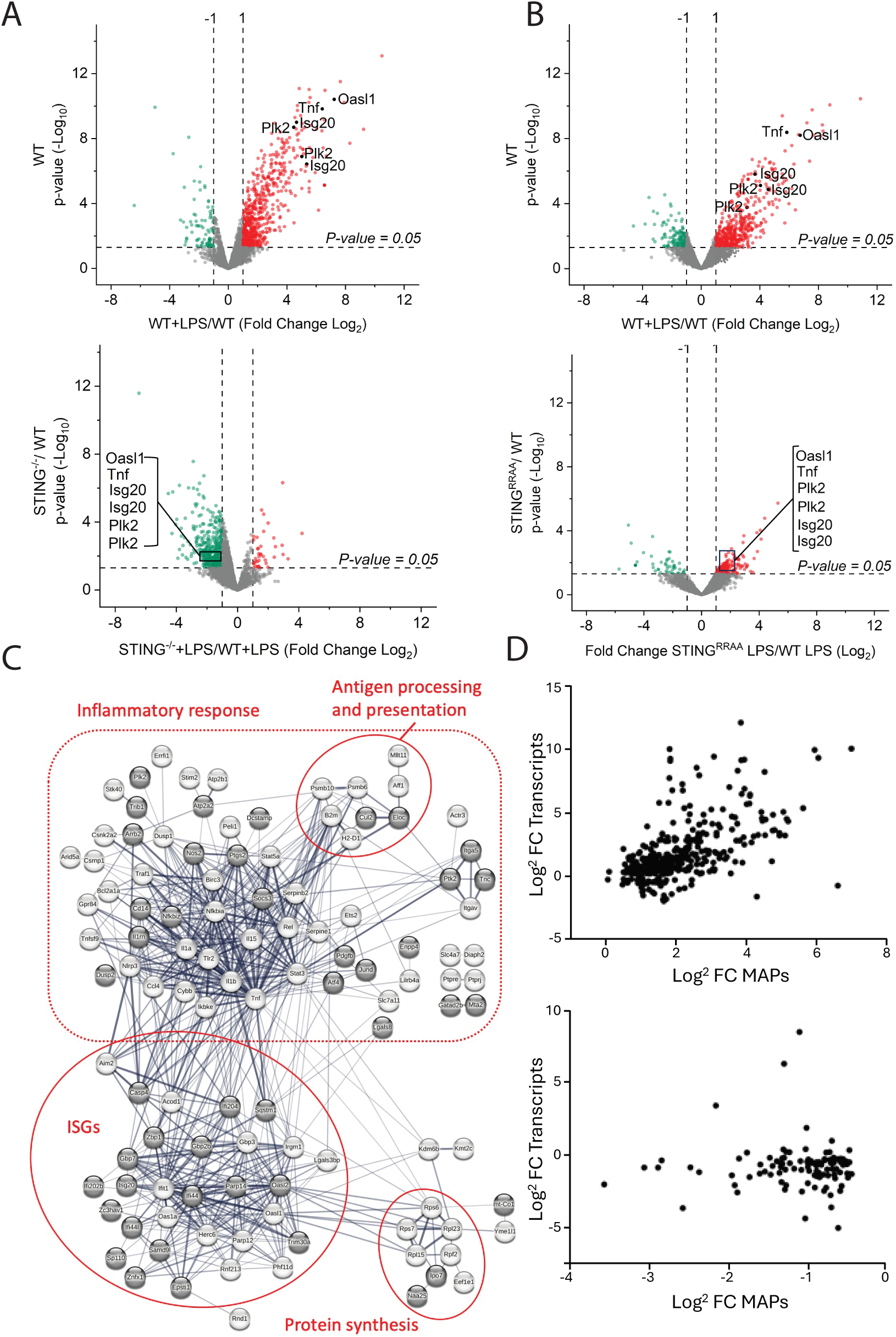
STING is a key regulator of antigen presentation. **A**) Volcano plot showing the effect of LPS on the abundance of the MAPs displayed at the surface of WT RAW cells (top). The bottom plot shows how the loss of STING (STING^-/-^) affects the abundance of MAPs displayed at the surface of RAW cells (STING^-/-^ +LPS/WT+LPS). **B**) Volcano plot showing the effect of LPS on the abundance of MAPs displayed at the surface of WT RAW cells (top) and how the loss of the UPR motif in STING^RRAA^ cells affects the abundance of MAPs displayed at the cell surface (STING^RRAA^ +LPS/WT +LPS). **C**) STRING interactome of the source proteins that had peptides presented in high amounts (all nodes > 5-fold) in response to LPS (WT +LPS/WT untreated). Proteins in black are the source of peptides whose presentation is down-regulated by at least 2-fold in response to LPS in STING^-/-^ cells (STING^-/-^ +LPS/WT +LPS). The edges indicate functional and physical protein associations. The line thickness indicates the strength of the data support. The assignment in the red circles and box was based on a literature survey. **D**) Top: Transcriptomics analyses showing the correlation between the increase in the presentation of peptides and the expression of the mRNA of their source proteins in response to LPS. Bottom: Transcriptomics analyses of the expression of the mRNA of source proteins that had peptides downregulated in response to LPS (more abundant on WT RAW cells before LPS treatment).

Altogether, these data suggest that STING acts as a rheostat to attenuate PERK activation and the magnitude of protein synthesis inhibition, enabling the expression of XBP1s and MitAP following TLR4 activation in WT RAW macrophages. In STING^RRAA^ cells the UPR motif is no longer there to dampen PERK activity and the pathway leading to translational shutoff. Accordingly, we reasoned that pharmacological inhibition of PERK using the specific inhibitor GSK2606414, prior to LPS treatment, would limit the extent of the translational shutoff induced by LPS and rescue MitAP in STING^RRAA^ cells. Indeed, the translational shutoff induced by LPS was limited by the pre-treatment with the PERK inhibitor (Fig. 4F) and MitAP was rescued (Fig. 4G).

The functional interaction between STING and PERK suggests that these proteins regulate MitAP from the endoplasmic reticulum (ER), where both are localized. Consistent with this model, Western blot analysis revealed that STING was neither phosphorylated nor degraded following up to 6 h of LPS treatment in WT RAW cells (Fig. 4H). This indicates that, under these conditions, STING does not traffic to the Golgi—where phosphorylation occurs—nor to lysosomes, where it is degraded. In contrast, direct activation of STING with the agonist DMXAA induced rapid phosphorylation and degradation of the protein (Fig. 4H).

To directly assess STING localization, we examined WT RAW cells stably expressing STING–GFP. LPS treatment failed to induce STING trafficking to the Golgi, as no colocalization with the Golgi marker GM130 was detected after 2 h (Fig. 4I), and even up to 6 h post-LPS (not shown). Importantly, LPS robustly induced MitAP under these conditions (Supp. Fig. 8A), indicating that STING supports this pathway while remaining localized to the ER. To exclude the possibility that the GFP tag interfered with STING trafficking, cells were stimulated with DMXAA. Under these conditions, STING–GFP rapidly translocated to the Golgi within 30–60 min and subsequently exited to post-Golgi vesicular compartments by 120 min (Fig. 4I). Together, these findings support the conclusion that STING regulates MitAP from the ER rather than through its canonical trafficking pathway.

We next examined whether the “UPR motif” of STING is required for regulating its exit from the ER. To this end, we stably expressed GFP-tagged STING variants harboring either the RRAA or RRDD substitutions within the “UPR motif”, which are known to differentially affect IFN-β signaling ^30^. These mutations had distinct effects on STING trafficking. In contrast to WT STING, LPS stimulation promoted the translocation of STING^RRAA^ to the Golgi (Supp. Fig. 8B), indicating that the UPR motif is required to retain STING within the ER during inflammatory signaling. Conversely, the STING^RRDD^ mutant failed to translocate to the Golgi in response to both LPS and the potent agonist DMXAA (Supp. Fig. 8C), demonstrating that the “UPR motif” also governs responsiveness to canonical STING activation.

Consistent with these observations, robust MitAP activation was observed in WT cells, in which STING remained ER-localized, whereas enforced exit of STING from the ER in STING^RRAA^ cells correlated with a markedly attenuated MitAP response (Fig. 4B). To directly test this relationship, WT RAW cells were pretreated with DMXAA for 60 min to drive STING export from the ER prior to LPS stimulation. Under these conditions, MitAP activation was strongly reduced compared with LPS treatment alone (Supp. Fig. 8D). Collectively, these data demonstrate that retention of STING within the ER is essential for efficient MitAP induction during inflammatory signaling.

Finally, our finding that STING regulates MitAP from the ER in a non-phosphorylated state suggests that this activity occurs independently of its canonical interaction with TBK1 at the Golgi, where STING phosphorylation normally occurs. This raises the question of how TBK1 contributes to MitAP (see Figs. 2D and E), given that its role cannot be readily explained by Golgi-associated STING signaling alone. In addition to its interaction with STING at the Golgi, TBK1 has been reported to function in mitophagy at mitochondria ^41^) and within endophagosomal compartments to regulate phagosome maturation ^42^This is particularly relevant in the present context, as both mitochondria and endo-lysosomal compartments have been implicated at early stages of the MitAP pathway (Figs. 3F and G; ^8^). We found that pharmacological inhibition of TBK1 with BX795 strongly suppressed the cellular response to LPS, as evidenced by the markedly reduced production of both ATF4 and XBP1s (Supp. Fig. 8E). These observations are consistent with a model in which TBK1 functions at an early step along the MitAP pathway, upstream of the ER, where ER-resident UPR sensors PERK and IRE1 drive ATF4 and XBP1s production.

### XBP1s is required for MitAP activation

While MitAP was inhibited in response to LPS in STING^RRAA^ cells (Fig. 4B), this presentation pathway was efficiently activated when these cells were treated with TG (Fig. 5A and Supp. Fig. 4B). Interestingly, in contrast to the results with LPS, the balance between ATF4 and XBP1s expression was back to the level of the WT cells when the UPR was induced using TG (Fig. 5B). These data further emphasized that MitAP is activated when XBP1s is expressed. To formally show a link between XBP1s and MitAP, we first inhibited IRE1, the UPR sensor responsible for XBP1s production, with the inhibitor Kira6 before LPS treatment and observed a dose-dependent decrease in MitAP (Fig. 5C). Inhibitors for the two other UPR sensors; Ceapin-A7 (ATF6 inhibitor) and GSK2606414 (PERK inhibitor) had no inhibitory effects on this pathway (Supp. Fig. 9). Finally, we generated XBP1^-/-^ RAW cells and observed that the loss of this protein did not impair IRE1 autophosphorylation (an upstream step in the UPR pathway) (Fig. 5D) but significantly inhibited MitAP in response to LPS (Fig. 5E).

### STING regulates MitAP in APCs to drive the activation and proliferation of mitochondria-specific PD-related CD8^+^ T cells

Two recent studies highlighted the significance of MitAP and CD8^+^ T cell activation in PD mouse models. The first study demonstrated that the motor symptoms in *Pink1^-/-^* mice, induced upon the activation of MitAP following gut infection with Gram-negative bacteria ^9^, were prevented by depleting the CD8^+^ T cell pool before infection ^10^. The second study showed that the adoptive transfer of cytotoxic CD8^+^ T cells from transgenic 2C mice, which express a TCR specific for an OGDH peptide presented via the MitAP pathway (Supp. Fig. 4), was sufficient to damage dopaminergic neurons and induce motor impairments. These impairments were reversible with L-DOPA treatment ^11^. To further explore whether STING-mediated regulation of MitAP in APCs affects CD8^+^ T cell activation, we conducted the following experiment (Fig. 6A): Purified 2C CD8^+^ T cells were incubated with PFA-fixed RAW cells that were pre-stimulated or not with LPS. T-cell activation and proliferation were subsequently assessed. Co-incubation with LPS-stimulated WT RAW cells resulted in a significant upregulation of key activation markers, CD25, CD44, and CD69 (Fig. 6B), alongside a robust proliferation (Fig. 6C). In contrast, when 2C T cells were exposed to LPS-treated RAW cells deficient in STING (STING ^-/-^) or possessing a defective STING UPR motif (STING^RRAA^), neither activation marker expression nor cell proliferation was observed (Fig. 6B and C). This disparity highlighted the indispensable role of the STING pathway in APCs for driving MitAP-dependent activation and proliferation of primary CD8+ T cells in a PD-related context.

### STING regulates the presentation of self-antigens on MHC-I molecules in inflammatory conditions

Studying the presentation of self-peptides at the surface of APCs is limited by the few specific T cell hybridomas available. While the hybridomas used here allowed us to define key steps of the MitAP pathway, it was not possible to evaluate how the presentation of self-peptides more broadly is affected during the inflammatory response, and the extent to which the STING-UPR pathway regulates this process. To overcome this limitation, we employed a high-sensitivity mass spectrometry-based immunopeptidomics approach to identify peptides presented on MHC-I at the surface of WT and the STING-modified RAW cells in response to LPS. Due to the large number of cells required and the complexity of the MHC-I immunoprecipitation protocols, experiments were performed independently for the STING^⁻/⁻^ and STING^RRAA^ cells. We first compared how TLR4 activation following a 6h LPS treatment affected the repertoire of peptides presented at the surface of WT RAW macrophages and STING^-/-^ cells. A total of 2692 peptides were identified in these experiments (Supp. Table 1). The analyses were performed on a total of 6 replicates per condition. Pairwise Pearson correlation coefficient analyses of peptide intensities showed a high to very high reproducibility between replicates (Supp. Fig. 10). In the WT control cells, LPS treatment was found to have a significant impact on antigen presentation, with a large group of self-peptides increased by more than 2-fold (Fig. 7A, upper panel, red dots). A smaller subset showed a decrease by the same threshold (Fig. 7A, upper panel, green dots). These data indicate that a rapid remodeling of the immunopeptidome occurs during the inflammatory response, resulting in the presentation of a new set of peptides, including peptides derived from proteins associated with the inflammatory response (e.g., TNF and PLK2) and interferon-stimulated genes (ISGs), such as Isg20 and Oasl1. Indeed, Gene Ontology (GO) enrichment analyses indicated that the upregulated peptides were predominantly associated with immune-related functions. On the other hand, the downregulated peptides were related to a variety of non-immune cellular functions (Supp. Fig. 11).

The loss of STING in the STING^-/-^ cells had a profound effect on the immunopeptidome presented in response to LPS. Indeed, comparative analyses of the peptide abundance ratios (STING^-/-^ +LPS/ WT +LPS) revealed that the presentation of most of the peptides upregulated in WT cells exhibited markedly reduced presentation in STING^-/-^ cells (Fig. 7A, lower panel, green dots), including peptides derived from source proteins associated with inflammation. To dissect whether this effect was mediated through the IFN-β or UPR arms of STING signaling, we turned to the STING^RRAA^ mutant, which retains the ability to induce IFN-β but lacks UPR modulation. In these experiments, 3,025 peptides were identified (Supp. Table 1). Interestingly, the immunopeptidome of STING^RRAA^ cells treated with LPS showed no reduction in the presentation of the LPS-induced peptides upregulated in WT cells; in fact, several of these peptides were further increased (Fig. 8B, lower panel, red dots), likely due to intact IFN-β signaling (see Fig. 4A). However, a subset of peptides upregulated in WT cells was significantly decreased in STING^RRAA^ cells (Fig. 7B, lower panel, green dots), suggesting that they are presented via a distinct pathway, unrelated to IFN-β signaling but requiring the modulation of the UPR by STING (see Supp. Table 2).

Remarkably, STRING network analyses focusing on peptides upregulated by more than 5-fold in WT cells after LPS treatment (all nodes in Fig. 7C) identified two main groups of highly interacting proteins as the source of these peptides. The first consisted predominantly of proteins associated with the inflammatory response, such as NF-kappa B inhibitor alpha (NFKBIA), TNF, Casp4, NLRP3, and IL1α and β. The second group of source proteins contained ISGs, a type of proteins expressed in response to type I interferon, playing key roles in antiviral defense, antiproliferative activities, and the stimulation of adaptive immunity ^43^. This non-random enrichment of inflammatory and ISG-associated peptides underscored how the innate and adaptive immune responses are closely coordinated during stress; LPS rapidly inducing the presentation of peptides from immune-related proteins. Because most of the peptides upregulated in response to LPS came from proteins associated with inflammation, we concluded that the immunopeptidome reflects the cell’s recent activity; peptides displayed on MHC-I molecules acting as molecular snapshots of intracellular events. This specific display likely plays a role in immunosurveillance mechanisms, enabling the immune system to monitor the state of cellular homeostasis in real-time. The effect of the loss of STING on the presentation of self-antigens is further illustrated in Fig. 7C. The black circles represent the source proteins whose peptide presentation was inhibited by more than 2-fold in response to LPS in STING^-/-^cells, compared to WT controls (STING^-/-^ +LPS/WT +LPS). These data highlight the fundamental role played by STING in shaping the immunopeptidome during inflammation. In contrast to STING^-/-^ cells, no noticeable inhibition was observed in the presentation of the peptides associated with the inflammatory response and ISGs after the loss of the “UPR motif” in STING RRAA cells (Supp. Fig. 12), possibly because IFN-β is still produced in these cells in response to LPS (Fig. 4A).

Finally, as discussed above, we hypothesized that the peptides presented in response to LPS are a display of the current state of cell activity, suggesting that they originate from proteins recently synthesized. To determine whether this could be the case, we integrated and analyzed immunopeptidomics and transcriptomics data from WT cells after a 6h LPS treatment, reasoning that the mRNA transcripts of the source proteins providing peptides should also be upregulated by LPS. A strong correlation was observed between the peptides upregulated by LPS and the increase of the mRNA transcripts associated with their source proteins (r=0.60, Fig. 7D). Interestingly, the correlation value for the peptides downregulated following LPS treatment (present at the surface of resting cells before the LPS treatment and replaced by other peptides during the LPS-induced inflammatory response) was very low (r=0.4), suggesting that those were unlikely to be derived from proteins recently synthesized.

## Discussion

The hallmark PD motor symptoms associated with dopaminergic neuron cell death are preceded by a prodromal period with no apparent neuron loss that can last for up to 20 years ^2,3^. The molecular mechanisms responsible for the onset and progression of the disease during that period are poorly understood. While aging is the primary risk factor for PD, recent studies suggest that inflammation in peripheral organs, such as the gut, may contribute to disease risk ^44–46^. In addition to aging and inflammation, emerging evidence points to a role for adaptive immunity in PD ^4^. In that context, we showed previously that the presentation of certain peptides from proteins present in the mitochondrial matrix (MitAP) is regulated by PD-related proteins ^8^. Furthermore, activation of the MitAP pathway after gut infection with Gram-negative bacteria in *Pink1^-/-^*mice led to the emergence of severe motor impairments, fully reversible by L-DOPA treatment ^9^. Here, we delineate the molecular pathways regulating MitAP and, more broadly, the presentation of self-antigens on MHC-I molecules during the inflammatory response, downstream of TLR4 activation.

In response to LPS, MitAP is engaged by a non-canonical pathway involving the TRAM/TRIF arm of TLR4, independently from IRF3. Instead, MitAP requires TBK1 and one of its interacting partners, STING. Engagement of the cGAS-STING pathway in response to LPS appears to be driven by DNA sensing in the cytoplasm. Although we have not formally identified the source of DNA responsible for cGAS activation by LPS during MitAP, our results point to a role for mtDNA. Indeed, we observed that LPS treatment triggers the release of mtDNA in small vesicular structures, similar to MDVs shown to participate in the transfer of mtDNA to the cytosol ^26,47^. Interestingly, MitAP is inhibited in cells where the expression of SNX9 is decreased^8^, a protein involved in MDVs formation^47^, further linking mtDNA to the presentation of mitochondrial antigens. The increase in cGAMP production in the cytosol in response to LPS confirms that cGAS is activated upon TLR4 engagement in our system. The mechanisms leading to the leakage of mtDNA from MDVs to the cytosol have not been addressed here. However, while studying the role of MAPL, a SUMO-ligase present on the mitochondrial membrane, in cell death, we showed that mtDNA is transferred from MDVs to the lumen of lysosomes, where gasdermin-pores assemble during inflammation, releasing mtDNA in the cytosol and the initiation of pyroptosis, in a STING-dependent manner ^26^.

Our data showed that neither IRF3 nor NF-kB, two effectors involved in innate immunity downstream of STING, are essential for MitAP, indicating that STING regulates MitAP through another pathway. The recent finding that STING displays a motif enabling its functional interaction with the UPR ^30^ led us to investigate whether STING may play a role in MitAP by regulating this stress response. Key differences in how the UPR is induced in WT cells and in cells with a dysfunctional STING UPR motif provided insight into how this protein plays a role in MitAP and the presentation of self-peptides. In WT cells, LPS stimulation induced the expression of both ATF4 and XBP1s, as previously reported ^36,48,49^. However, two hallmark features of a canonical UPR—eIF2α phosphorylation and the resulting translational shutoff—were not observed. The absence of translational inhibition likely explains why XBP1s, whose synthesis is sensitive to global translation rates, is produced in WT RAW cells. In contrast, disruption of the STING “UPR motif” in STING^RRAA^ cells resulted in strong eIF2α phosphorylation and sustained translational inhibition following LPS treatment, leading to a marked reduction in XBP1s expression. Based on these observations, we propose that STING limits PERK activation in WT cells to dampen the magnitude of the UPR during inflammation, thereby permitting XBP1s production, a protein we showed to be required for MitAP. Attenuation of PERK activity and the ensuing stress responses could prevent premature cell death and prolong macrophage survival, as previously shown for dendritic cells ^50^

The mechanisms by which STING interacts with and modulates the UPR along the MitAP pathway is unknown. Upon activation, STING was shown to translocate from the ER to the trans-Golgi network (TGN), where it is phosphorylated by TBK1 and contributes to NF-κB and IRF3 activation ^51–53^. The observation that neither of these effectors are required for MitAP suggested that STING plays a role along this presentation pathway elsewhere. Instead, a key phenomenon observed in both the STING KO and RRAA cells is the delayed and decreased production of ATF4 induced in response to LPS, suggesting that STING restrains PERK activity. Given its proximity to UPR sensors in the ER, STING may functionally interact with the UPR in this compartment. Both our immunofluorescence and Western blot data indicate that STING modulates the UPR and MitAP while remaining in the ER in a non-phosphorylated form. This step appears to be regulated, at least in part, by the “UPR motif,” as evidenced by our observation that, unlike its WT counterpart, STING^RRAA^ translocates to the Golgi after LPS treatment, whereas STING^RRDD^ remains confined to the ER even when cells are stimulated with the potent STING agonist DMXAA. Consistent with this idea, mutations in STING have been shown to affect its trafficking. Notably, the N154S mutation associated with the STING-associated vasculopathy with onset in infancy (SAVI) was shown to cause a premature exit of STING from the ER and its accumulation in the Golgi ^54^. Interestingly, the STING^RRDD^ double substitution within the UPR motif prevented this exit from the ER ^30^. The differential effects ofof the RRAA and RRDD mutation on STING retention/exit from the ER may be related to the finding that, unlike RRAA, the ER-retained RRDD mutant affects the IFN-β pathway (IFIT1 expression) ^30^.

Although we did not detect a significant traffic of STING from the ER to the Golgi and lysosomes in response to LPS by immunofluorescence, nor noticeable phosphorylation and degradation by Western blot, we cannot rule out that STING may also regulate certain aspects of the MitAP pathway and self-antigen presentation from these compartments. Indeed, STING may also function in lysosomes where it has been shown to act as a proton channel ^55^. Recent work in human cells demonstrated that STING participates in lysosomal quality control and repair through this activity ^56^. Consistent with this, we previously showed that processing and presentation of mitochondria-derived peptides, which is initiated in lysosomes, is altered when lysosomal function is impaired ^8^. A deeper understanding of how STING trafficking from the ER to the TGN and lysosomes shapes its functions will improve our ability to manipulate STING–UPR signaling and develop therapeutic strategies for diseases associated with dysregulation of these pathways ^57,58^.

Altogether, our findings support the concept that MitAP is a regulated pathway responsive to inflammatory stimuli, rather than a stochastic phenomenon. While the precise identity of the mitochondrial-derived peptidome presented by MitAP remains to be fully characterized, the pathway defined in the handling of both gB and OGDH peptides engages common molecular checkpoints that confer both specificity and reproducibility. Importantly, the OGDH-derived peptide is functionally relevant, as evidenced by its ability to activate and drive the proliferation of CD8^+^ T cells. Furthermore, CD8+ T cells specific for the OGDH peptide presented via the MitAP pathway are directly responsible for the emergence of severe PD-like motor impairment, reversible by L-DOPA treatment following adoptive transfer in mice ^11^.

In addition to MitAP, the STING-UPR pathway affects the presentation of self-peptides more broadly (see Fig. 8). STING appears to exert this effect through two complementary mechanisms: regulation of the IFN-β pathway and modulation of the UPR. In WT cells, activation of the IFN-β pathway by LPS enables the presentation of a series of peptides originating from proteins associated with the inflammatory response, including several ISGs. We argued that since most of these proteins are not present in resting cells, these peptides originate from proteins synthesized de novo, following the sensing of “danger” signals during inflammation. This rapid crosstalk between the innate and adaptive immune responses may represent a component of immune-surveillance mechanisms monitoring cell activity in “real-time”. The preferential presentation of peptides from proteins expressed as a rapid response to a danger cue aligns with the DRiP hypothesis, first proposed by the group of Jon Yewdell ^59^, which posits that prematurely terminated polypeptides and misfolded proteins serve as a primary source of endogenous peptides presented on MHC-I molecules. Further studies, focusing on the identification of newly translated mRNA transcripts using RiboSeq, will, however, be required to identify the presented peptides as DRiPs. These investigations will be crucial for defining the contribution of DRiPs to antigen presentation during inflammation and for assessing their role in immune surveillance and autoimmunity.

In STING^-/-^ cells, where the control over both the IFN-β and UPR pathways is affected, the presentation of several of the inflammatory and ISG-derived peptides, as well as MitAP, is inhibited. Remarkably, in STING^RRAA^ cells, where the IFN-β pathway is preserved but the UPR modulation is lost, MitAP is inhibited, while the presentation of peptides from proteins linked to the inflammatory response and ISGs is maintained. Thus, two distinct pathways appear to be regulated by STING. The first one, driven by the IFN-β pathway, enables the presentation of a set of peptides related to the inflammatory response that may contribute to immunosurveillance. The second one, regulated by the magnitude of the UPR during inflammation, appears to be responsible for the presentation of a more restricted set of peptides, including, the gB and OGDH peptides of the MitAP pathway, and a set of peptides from a variety of proteins (Supp. Table 2). Whether this pathway plays a role in immune surveillance, or the initiation of autoimmune mechanisms remains to be established.

The involvement of STING and the UPR in antigen presentation has interesting implications for the management of inflammatory and autoimmune diseases, as well as cancer and degenerative diseases linked to aging. Indeed, our data indicate that targeting the STING-UPR axis may have potential therapeutic value not only in regulating the inflammatory response but also in fine-tuning antigen presentation, thereby downregulating this process in autoimmune diseases and promoting it in certain cancers that evade immune system recognition, as suggested recently ^60^. Modulating the cGAS-STING pathway has already shown promise in preclinical models of autoimmunity and cancer, with small-molecule inhibitors and agonists currently under development ^61,62^. Similarly, targeting UPR components offers another avenue for intervention. Compounds modulating IRE1α or restoring XBP1s expression could preserve MitAP while minimizing the risk of global protein synthesis inhibition. A better understanding of the molecular mechanisms that enable the TLR4-STING-UPR pathway to regulate innate and adaptive immune responses would facilitate the development of precision approaches that can mitigate inflammation and autoimmunity with reduced off-target effects.

By defining the cGAS-STING-UPR axis as a central regulator of MitAP, our study bridges critical gaps in understanding the interplay between inflammation, antigen presentation, and autoimmunity in a PD-related pathway. This integrated model not only advances fundamental knowledge but also identifies actionable targets for therapeutic development. Future efforts should focus on translating these findings into clinical applications, including immunopeptidomics-guided diagnostics and STING-UPR-directed therapies, to address the unmet needs of PD and related disorders.

### Limitations of the Study

The work presented here dissects a pathway initiated at the cell surface by TLR4 engagement that culminates in the presentation of self-antigens, including mitochondrial matrix–derived antigens presented through the MitAP pathway. Several key questions remain unresolved. These include how TLR4 signaling promotes the formation of mtDNA-containing MDVs; whether mtDNA is released from lysosomes following MDV–lysosome fusion, as recently described^26^; and which events downstream of XBP1s lead to MitAP activation and self-antigen presentation. Although we demonstrate that STING acts from the ER to activate MitAP, the precise mechanisms by which STING regulates the UPR remain to be defined. For example, considering that cGAS is required for MitAP, it will be important to determine the precise role of its product, cGAMP, in the regulation of STING function within the ER. It is also imperative to determine whether the “UPR motif” of STING directly interacts with PERK to modulate the UPR. The mechanism by which TBK1 regulates MitAP, independently of STING phosphorylation, also remains to be elucidated. Finally, while MitAP has been linked to the initiation of autoimmune mechanisms and PD-like symptoms in a mouse model, its relevance in humans remains to be established. Progress in this area is currently limited by the scarcity of tools to study antigen presentation in human systems. The application of immunopeptidomics to human samples may therefore provide new opportunities to elucidate adaptive immune mechanisms in people with Parkinson’s disease.

## Resource Availability

### Lead contact

Requests for further information, resources, and reagents should be directed to and will be fulfilled by the lead contact, Michel Desjardins.

### Materials availability

Key lab materials used and generated in this study are listed in a Key Resource Table alongside their persistent identifiers at[DOI:10.5281/zenodo.19560905.].

### Data and code availability

The data, protocols used and generated in this study are listed in a Key Resource Table alongside their persistent identifiers at [10.5281/zenodo.19560905.].

Raw and processed transcriptomic data are available in GSE329088. Immunopeptidomics data have been deposited to the ProteomeXchange Consortium via the PRIDE partner repository (dataset identifier PXD059785). No code was generated for this study.

## Acknowledgements

The study is funded by the joint efforts of The Michael J. Fox Foundation for Parkinson’s Research (MJFF) and the Aligning Science Across Parkinson’s (ASAP) initiative. MJFF administers the grant ASAP 000525 on behalf of ASAP and itself. The Institute for Research in Immunology and Cancer (IRIC) receives infrastructure support from IRICoR, the Canadian Foundation for Innovation, and the Fonds de Recherche du Québec - Santé (FRQS). We thank the Biomedical Innovation Centre (CIB) and Dr. Nicolas Stifani for providing access to, and assistance with, the microscopy facility. This work was also supported by CIHR grants to ES (FDN-143332), SG (PJT-162406), and MD (169204). The authors thank Dr. Suzanne Pfeffer and Dr. Nathalie Labrecque for their constructive comments and discussions. We sincerely thank Dr. Lilia Rodriguez Moya for her valuable assistance in handling and proofreading the manuscript.

## Author contributions

**A.M.F.**: Conceptualization, Formal analysis, Investigation, Methodology, Supervision, Visualization, Writing-original draft, Writing-review & editing; **A.A.**: Investigation, Methodology, Formal analysis. **J.L.**: Formal analysis, Methodology, Data curation. **T.C.**: Investigation, Formal analysis. **M.N.E.**: Investigation, Methodology, Formal analysis. **C.H.P.**: Investigation, Formal analysis. **B.B.**: Investigation, Methodology. **E.B.**: Data curation. **Y.Z.X.**: Methodology. **M.I.**: Investigation. **G.A.D.**: Investigation. **E.O.A.**: Data curation. **S.L.**: Data curation, Supervision. **E.C.**: Conceptualization. **E.S.**: Funding acquisition. Resources, Supervision, Writing-review & editing. **P.P.**: Conceptualization, Writing-review & editing. **S.G.**: Conceptualization, Funding acquisition, Supervision. **P.T.**: Conceptualization, Funding acquisition, Supervision, Writing-review & editing. **H.M.M.**: Conceptualization, Funding acquisition, Writing-review & editing. **M.D.**: Conceptualization, Formal analysis, Funding acquisition, Investigation, Project administration, Supervision, Visualization, Writing-original draft, Writing-review & editing

## STAR★Methods

### Key resources table

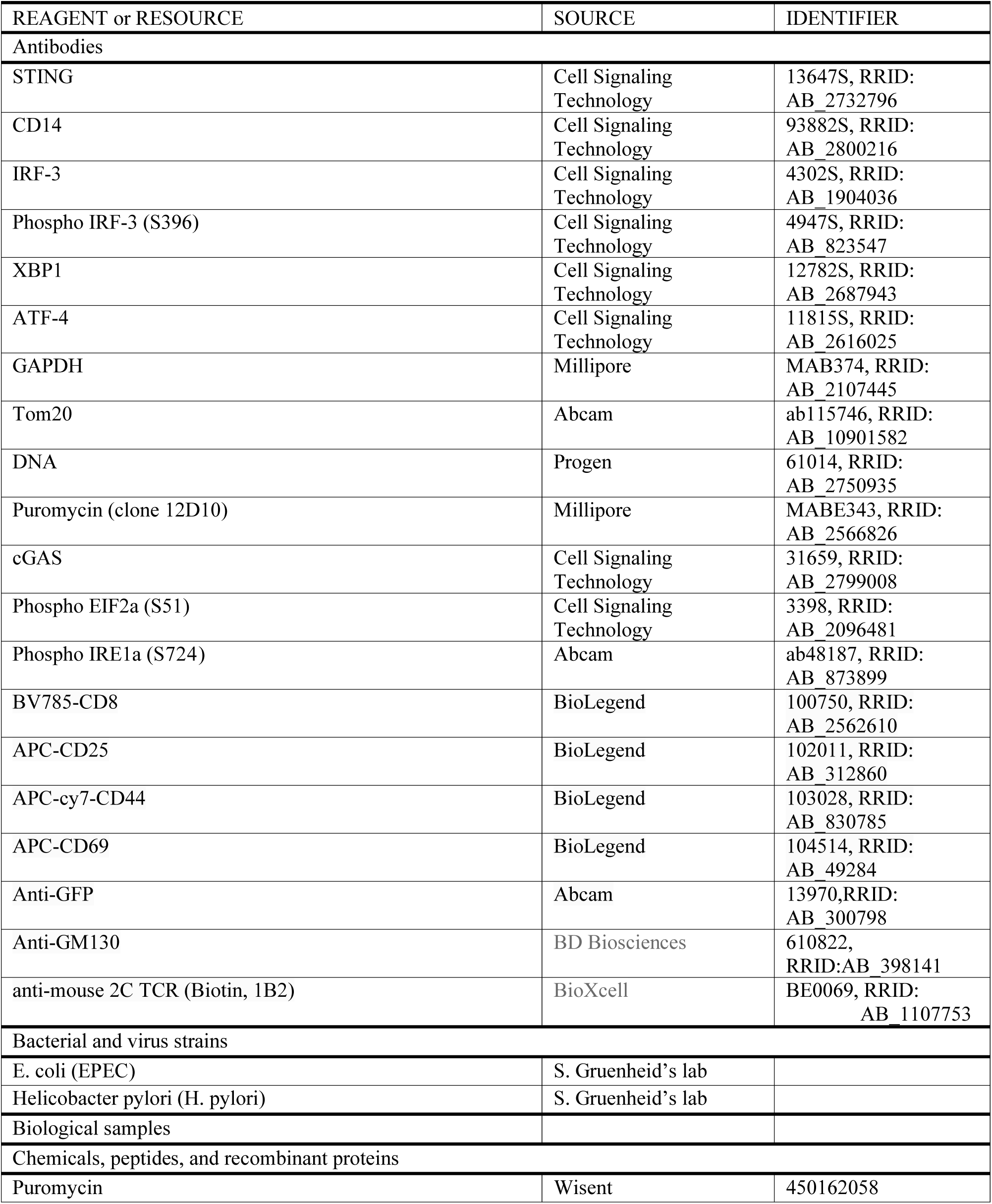
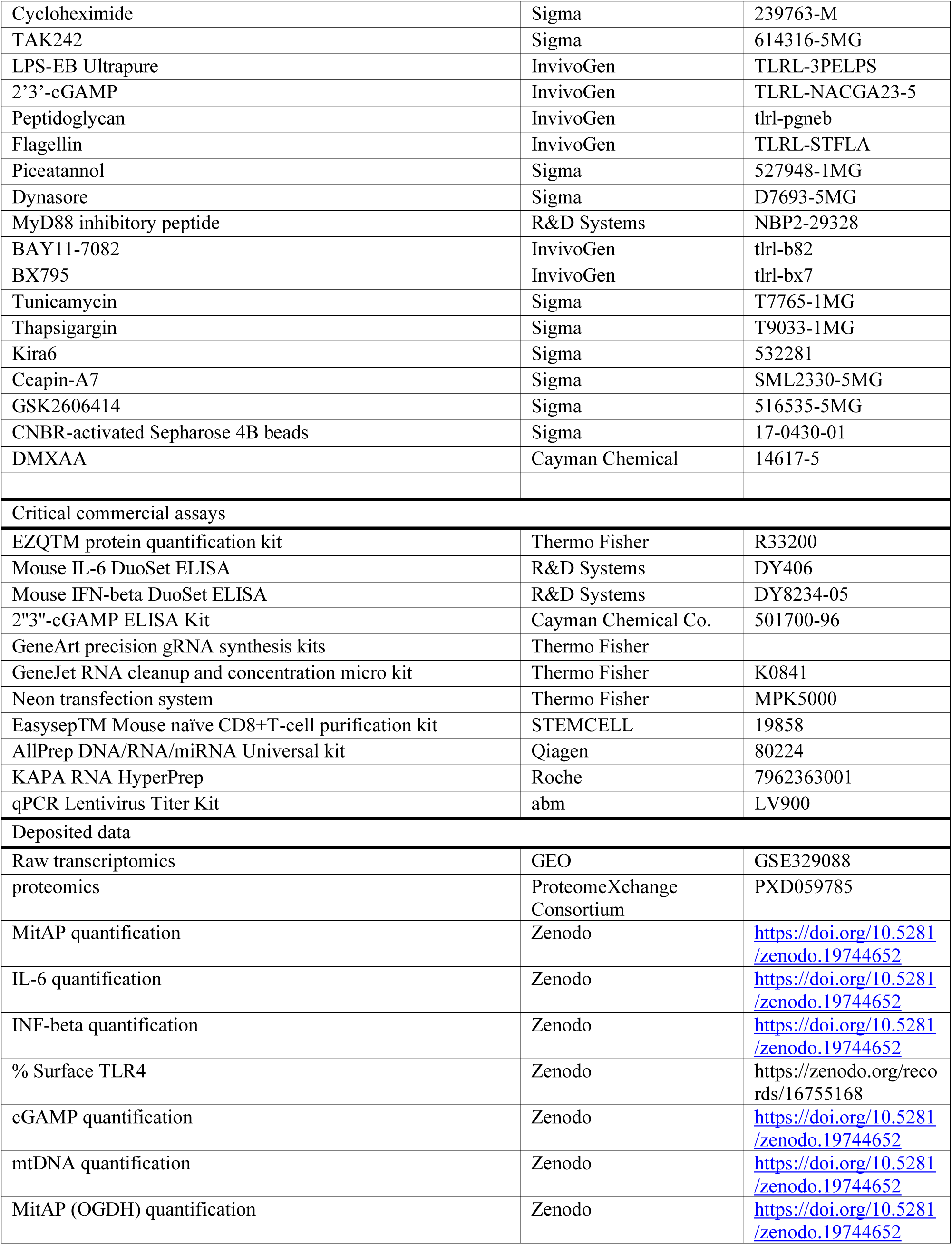
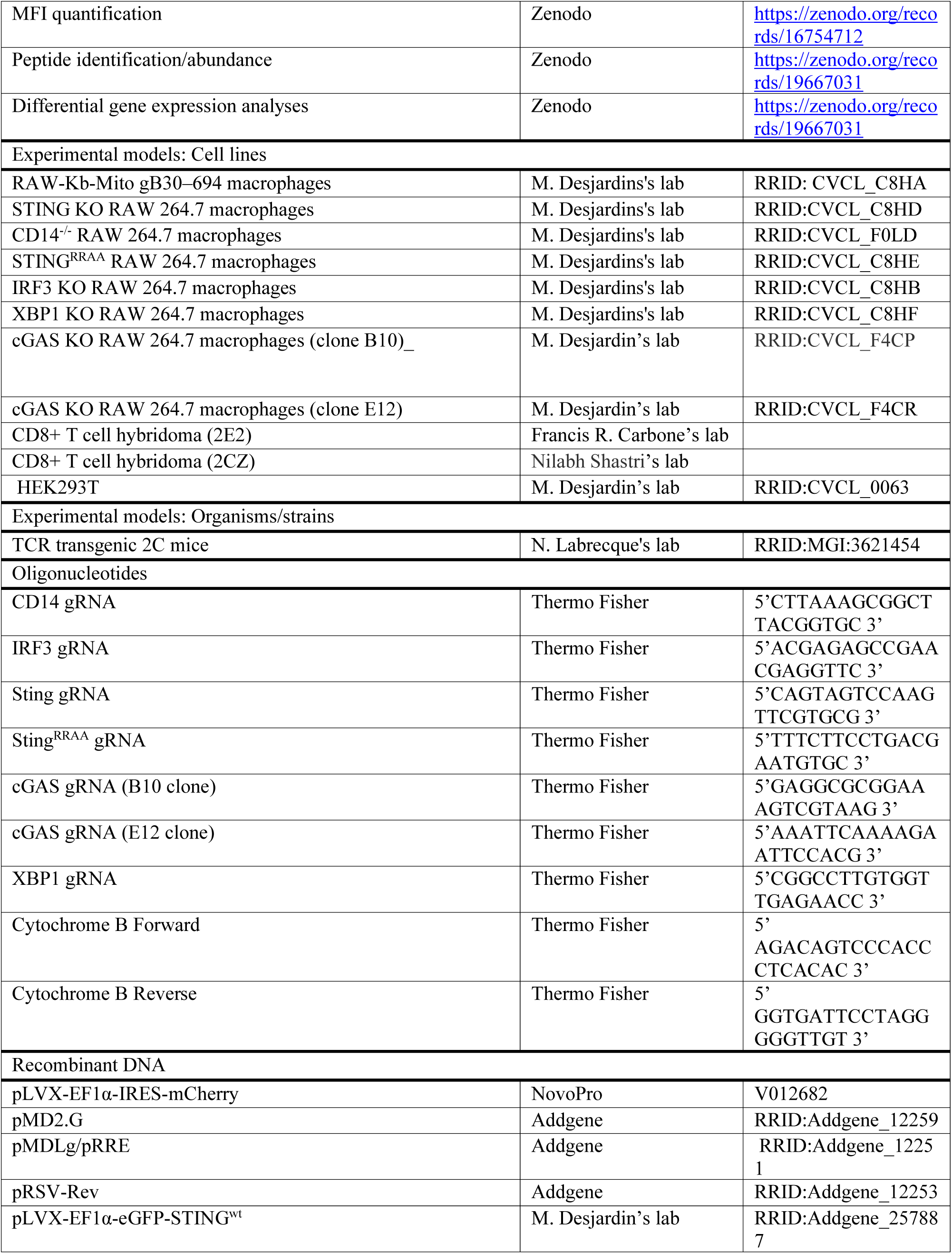
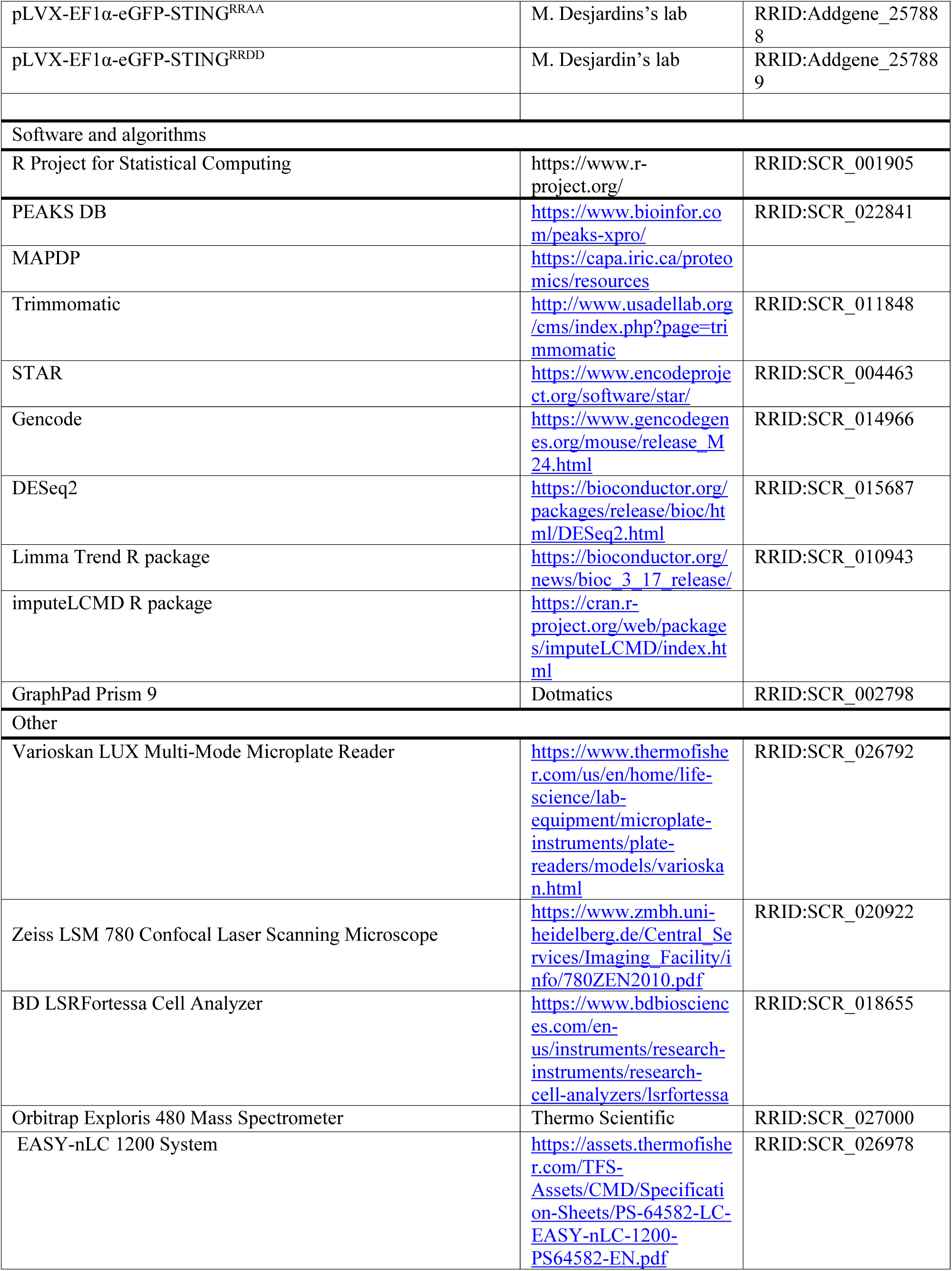
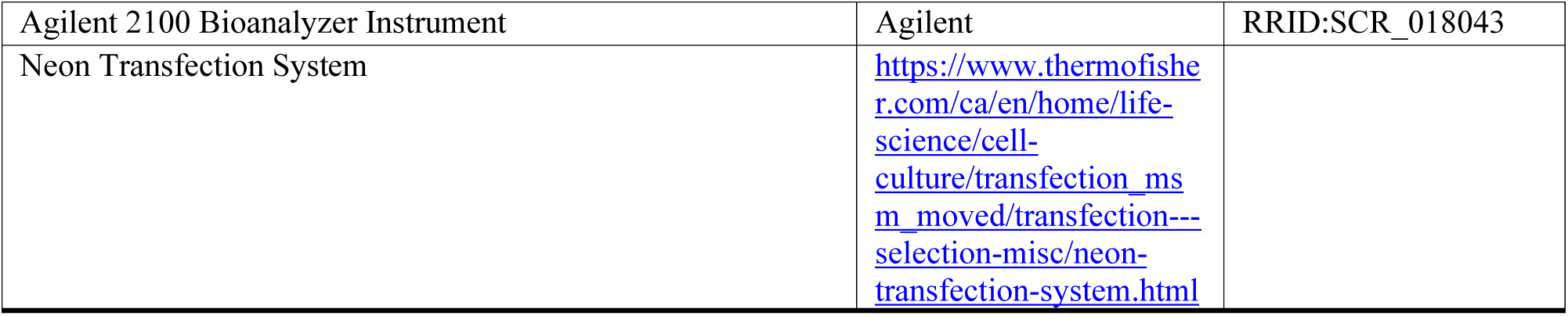

### EXPERIMENTAL MODEL DETAILS

#### Animal handling and cell cultures

The TCR transgenic 2C T cells were isolated from male mice in strict accordance with good animal practice as defined by the Canadian Council on Animal Care and following protocols approved by the Comité de déontologie animale of the Université de Montréal. This study was performed using only male mice, which limits the potential influence of gender. The RAW 264.7 macrophage cell line was obtained from ATCC (TIB-71) and modified to express H-2K^b^ and the glycoprotein B (gB) from HSV-1 in the mitochondria matrix as previously described ^8^. Cells were cultured in DMEM with 10% (v/v) fetal calf serum (FCS), penicillin (100 U ml^−1^) and streptomycin (100 μg ml^−1^). The β-galactosidase-inducible HSV gB/K^b^-restricted HSV- 2.3.2E2 CD8^+^ T cell hybridoma (2E2) was provided by Francis R. Carbone’s lab at the University of Melbourne ^63^. The OGDH/L^d^ and OGDH/K^b^-restricted 2CZ CD8T^+^ cell hybridoma was provided by Dr. Nilabh Shastri’s lab at the University of California at Berkeley ^64,65^. Hybridomas were maintained in RPMI-1640 medium supplemented with 5% (v/v) FCS, glutamine (2 mM), penicillin (100 U ml^−1^) and streptomycin (100 μg ml^−1^). All cell lines were regularly tested for Mycoplasma.

#### Bacterial Preparation and Cell Infection

*Helicobacter pylori* (PMSS1 strain) and *Escherichia coli* (EPEC) (E2348/69 strain) were grown overnight in LB broth at 37 °C with shaking. Cultures were then diluted in DMEM supplemented with 10% fetal bovine serum (FBS) to a final concentration of 1 × 10⁵ CFU/μL and used to infect cells at a multiplicity of infection (MOI) of 1. Heat-killed EPEC was generated by incubating the diluted EPEC culture at 95 °C for 10 minutes and subsequently applied to cells at a 1:1 ratio.

### METHOD DETAILS

#### Reagents and Antibodies

Thapsigargin (T9033-1MG), Tunicamycin (T7765-1MG), ceapin-A7 (SML2330-5MG), PERK Inhibitor I, GSK2606414 (516535-5MG), IRE1α Inhibitor IV, KIRA6 (532281), TAK-242 (614316-5MG), Dynasore hydrate (D7693-5MG), and Piceatannol (527948-1MG) were purchased from Sigma. BAY11-7082 (tlrl-b82), LPS-EB Ultrapure (TLRL-3PELPS), 2’3’-cGAMP (TLRL-NACGA23-5), PGN-EB (tlrl-pgneb), Fla-st (TLRL-STFLA), and BX795 (tlrl-bx7) were from InvivoGen. MyD88 Inhibitor Peptide (NBP2-29328) was from R&D systems. DMXAA (14617-5) was from Cayman chemical.Antibodies against STING (13647S, RRID: AB_2732796), CD14 (93882S, RRID: AB_2800216), IRF-3 (4302S, RRID: AB_1904036), Phospho-IRF-3 (4947S, RRID: AB_823547), XBP-1s (12782S, RRID: AB_2687943), and ATF-4 (11815S, RRID: AB_2616025) were purchased from Cell Signalling. Anti-GAPDH (MAB374, RRID: AB_2107445) was from Millipore. Anti-Tom20 (ab115746, RRID: AB_10901582) was from Abcam. Anti-DNA (61014, RRID: AB_2750935) was from PROGEN. Anti-GFP (13970,RRID: AB_300798) was from Abcam. Anti-GM130 (610822, RRID:AB_398141) was from BD Biosciences. The Anti-Puromycin clone 12D10 (MABE343, RRID: AB_2566826) antibody was obtained from Millipore. FACS antibodies targeting BV785-CD8 (100750, RRID: AB_2562610), APC-CD25 (102011, RRID: AB_312860), APC-cy7-CD44 (103028, RRID: AB_830785), and APC-CD69 (104514, RRID: AB_49284) were purchased from BioLegend. The Anti-1B2 (Biotin) antibody is available at BioXcell (Cat# BE0069, RRID:AB_1107753)

#### Western Blotting

Cells were lysed in Laemmli buffer, and the protein content was quantified using EZQ^TM^ protein quantification kit (Thermo Fisher). Equal protein amounts (40 μg) were loaded and separated on 4%–15% pre-cast gels (Bio-Rad) and transferred using a Trans-Blot turbo system (Bio-Rad). Membranes were blocked either in 5% milk TBS or 5%BSA in TBS for 30 min at room temperature and incubated with the primary antibody diluted in 5% BSA-TBS overnight at 4°C. The following day, the membrane was washed 3 times in Tris-buffered saline – 0.4 % Tween(20)(TBS-T), incubated with the secondary antibody (diluted in TBS-T plus 5% milk or 5% BSA) for 1 h at room temperature and washed 3 times in TBS-T. The membrane was then developed in Clarity Western ECL substrate (BioRad) and visualized using the ChemiDoc™ imaging system (Bio-Rad).

#### ELISA

Collected supernatants were assayed for IL-6 and IFN-β using mouse DuoSet kits (R&D Systems). Samples were diluted in reagent diluent and incubated in a 96-well plate pre-coated with either IL-6 or IFN-β capture antibodies for 2 h at room temperature. Plates were then washed three times with PBS plus 0.05% Tween 20 and incubated with the respective detection antibody for 2 h at room temperature, followed by three washes. Plates were finally incubated with streptavidin–HRP in the dark for 20 min, washed three times and developed with a 1:1 ratio of hydrogen peroxide and tetramethylbenzidine (Thermo Fisher). The optical density of the plates was read at 450 nm using a Varioskan microplate reader (Thermo Fisher). The 2’’3’’-cGAMP ELISA was performed using the 2’’3’’-cGAMP ELISA Kit (Cayman Chemical Co.) according to the manufacturer’s instructions.

#### Mitochondrial antigen-presentation assay

RAW cells were seeded at a density of 1×10⁶ cells per well in a 24-well plate one day before treatment. The following day, cells were treated either with LPS (1.5 μg/mL) or the specified inducers for 6 hours. For experiments involving inhibitors, cells were pre-treated with the inhibitor for 2 hours before LPS stimulation. Cells were transferred into a 96-well plate 1 hour before fixation. Cells were fixed with 1% paraformaldehyde (PFA) for 10 min at room temperature, washed 5 times with wash media (RPMI, 10% FBS and 0.1 M glycine) and incubated with 1 × 10^5^ 2E2 T cell hybridoma (ratio 3:1) for 16 h. β-galactosidase activity was then measured at 595 nm after the addition of CPRG substrate using a Varioskan microplate reader (Thermo Fisher). MitAP was then presented as fold induction over the untreated control. Since all the RAW cell types modified by CRISPR have their own WT control cells (CRISPR with an empty vector), there could be variations in the levels of MitAP activation that do not affect results within a particular modified cell type.

##### Immunofluorescence

RAW cells were fixed with 5% PFA for 15 min at 37 °C. The PFA was then quenched with 50 mM NH_4_Cl in PBS for 10 min at room temperature. Cells were permeabilized with 0.1% Triton X- 100 in PBS (v/v) for 10 min at room temperature and then blocked with blocking buffer (5% FBS in PBS) for 10 min at room temperature. Cells were incubated with the indicated primary antibodies for 16 h at 4°C. After three washes with PBS, cells were incubated with the appropriate secondary antibodies anti-rabbit-A488(A-11008, RRID: AB_143165) and anti-mouse-A568 (A-11004, RRID: AB_2534072), anti-Chicken-488 (A-11039, RRID:AB_142924) Life Technologies, 1:1000, for 1 h. After being washed and mounted with Prolong™ Antifade (Invitrogen), the slides were examined with a Zeiss LSM 780 or LSM 900 confocal microscope with Airyscan (Zeiss). For depletion of mtDNA, cells were treated with EtBr (200ng/ml) for 3 days, treated or not with LPS (1.5 μg/mL) for 6 h and then fixed and immunostained as described above.

#### CRISPR-Cas9 gene editing

The gRNAs for the generation of knock-out and knock-in) RAW cells were synthesized using GeneArt precision gRNA synthesis kits (Thermo Fisher) according to the manufacturer’s instructions. Before making gRNAs, 34-nucleotide forward and reverse target DNA oligonucleotides were designed using the CRISPR search and design tool (Thermo Fisher) and synthesized. The gRNA sequences are provided in the table below. The gRNA DNA templates were then PCR assembled, and gRNAs were synthesized by *in vitro* transcription. Then, gRNAs were purified (GeneJet RNA cleanup and concentration micro kit, ThermoScientific), and their concentrations were assessed. Before electroporation, Raw cells were scraped, washed with PBS, and counted. 1 x 10^5^ cells were then electroporated with 2μg Cas9 protein and 400 ng gRNA in 10 μl of resuspension buffer R (and 50 pmol of donor HDR templates in case of KI) using Neon transfection system (Thermo Fisher) according to the manufacturer’s instructions. After electroporation, cells were immediately transferred into prewarmed 1 ml of antibiotic-free growth medium in a well of a 12-well plate and cultured for 2 days. Cells were counted and serially diluted to 4 cells/ml. 200 μl of cell suspension was dispensed in each well of a 96-well plate. Plates were then incubated at 37°C in a 5% CO_2_ incubator for 10 days or until single clonal individual colonies can be observed under the microscope (5X magnification). Genomic DNA was isolated from single clones. The gRNA target region was amplified by PCR using AmpliTaq Gold 360 master mix (Thermo Fisher). PCR amplicons were sequenced using standard Sanger sequencing. Biallelic genomic Cas9 edits leading to frameshift mutation or HDR ODN insertions were confirmed by TIDE analysis ^66^. KO cells were further validated by Western blotting.

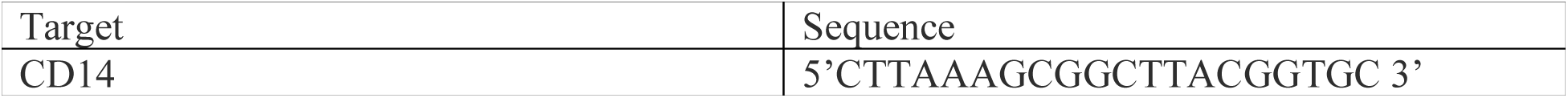
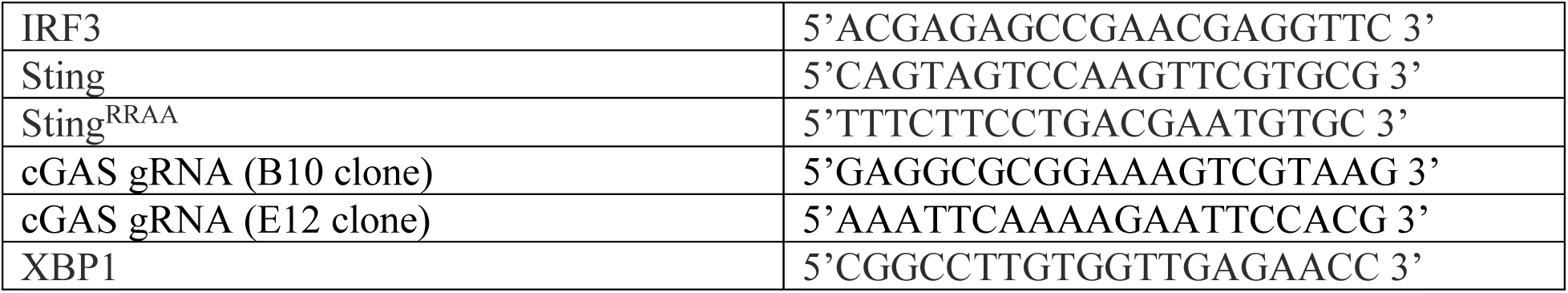

#### Stable cell line generation

Lentivirus were generated by co-transfecting HEK293T cells with pLVX-EF1α-IRES-mCherry vector (a kind gift from Ruth Slack, University of Ottawa) in which IRES-mCherry portion was replaced by PCR-amplified g-blocks (IDT) encoding GFP-STING^WT^, GFP-STING^RRAA^, or GFP-STING^RRDD^ sequences, together with the packaging plasmids pMD2.G (VSV-G), pMDLg/pRRE, and pRSV-Rev. Lentivirus titers were evaluated by qPCR kit (abm). Following 0.45 μm filtration, supernatants were concentrated (Lenti-X^TM^ concentrator, 631231, Takara) and added to RAW cells at a multiplicity of infection of at least 2. After 2 days, cells were passed and subsequently sorted based on GFP fluorescence by fluorescence-activated cell sorting.

#### Quantification of mtDNA by qPCR

RAW cells (4X10^6^) were plated in 6 well plates and stimulated with LPS (1.5 μg/mL) for the indicated time points. The cytosolic fraction was then collected, and mtDNA was isolated and quantified by qPCR using primers listed below, as previously described ^67^.

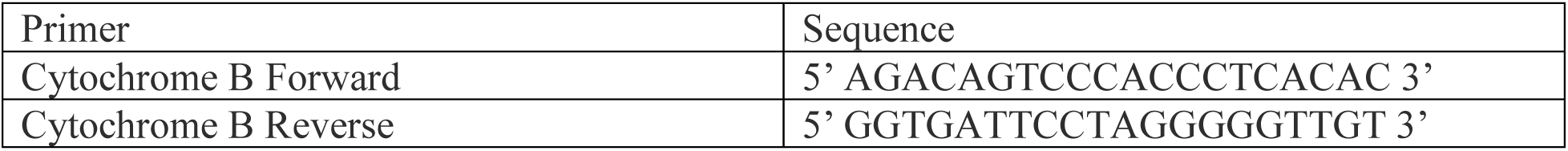

#### Flow cytometry

TLR4 internalization assessment was done as previously described ^68^. RAW cells (0.5X10^6^) were stimulated with LPS (1.5 μg/mL) for the indicated time points. Cells were then washed with 1 ml cold PBS. Live/dead cells were separated by ZombieAqua (BioLegend, #423102) and stained (30 min at 4°C) with PE anti-TLR4 (BioLegend; clone Sa15-21; 145404, RRID: AB_2561874) in blocking buffer (PBS with 2% rat serum). The stained cells were then washed with 1 ml cold PBS and resuspended in 200 ml PBS. Staining of the surface receptors was analyzed with BD FACSCanto II. The mean fluorescence intensity (MFI) of TLR4 from unstimulated or LPS-stimulated cells was recorded. The percentage of surface receptor staining at the indicated time points, which is the ratio of the MFI values measured from the stimulated cells to those measured from the unstimulated cells, was plotted to reflect the efficiency of receptor endocytosis.

For CD8+ T cell coculture experiments, naïve CD8+ T cells from the spleen of 2C TCR transgenic mice were purified using a Easysep^TM^ Mouse naïve CD8+T-cell purification kit (STEMCELL, BC, Canada). Purified CD8^+^ T cells were then labelled with cell trace violet (CTV) (Life Technologies, USA) (1μM for 10M cells), washed 2 times with PBS and resuspended in complete medium (RPMI 1640 supplemented with 2nM glutamine, penicillin/streptomycin, 10μM 2-mercaptoethanol, 0.1mM non-essential amino acids, 10mM HEPES, 1mM sodium pyruvate and 10% heat-inactivated fetal bovine serum). Different RAW cells were treated or not with LPS (1.5 μg/mL) for 6h, then fixed with 1% PFA for 10 min at room temperature, washed 3 times with wash media (RPMI, 10% FBS and 0.1 M glycine) then a final wash with complete medium. RAW and CD8+ T cells were incubated together at a 1:1 ratio. Loosely adherent cells were harvested on day 1 and day 3 post-incubation and stained with Zombie NIR^TM^viability dye (1:1000) for 20 min at 37°C, followed by anti-CD16/CD32 for another 20 min at 37°C. Samples were then stained for (CD8, CD25, CD44, CD69, anti-mouse 2C TCR antibody 1B2) and analyzed using BD LSRFortessa Cell Analyzer.

#### Nascent protein synthesis detection

Raw cells (2X10^6^) were treated with LPS (1.5 μg/mL) or LPS and cycloheximide (CHZ) (1μM) (Sigma) for the indicated time points. Cells were then labelled with puromycin (10μg/ml) for 10 min and washed three times with TBS. They were then lysed directly in Laemmli buffer, and the protein content was quantified using an EZQ^TM^ protein quantification kit (Thermo Fisher). An equal quantity of proteins was used for western blotting using an anti-puromycin antibody (12D10, Sigma).

For flow cytometry analysis, RAW cells (1 X10^6^) were pretreated with CHZ (1μM) for 10 minutes at 37°C as a negative control. Subsequently, cells were stimulated with LPS at a concentration of 1.5 μg/ml at the indicated time points, including the CHZ condition. Puromycin solution (10 μg/ml) (Wisent) was added, and cells were incubated for 30 minutes at 37°C and 5% CO2 (pulse). Following the pulse, cells were washed three times with 500 μl prewarmed DMEM and incubated for 1 hour at 37°C and 5% CO2 (chase) while maintaining treatments. Cells were then harvested and washed twice with cold FACS buffer (PBS+ 2% FBS), followed by a final wash with 1 ml cold PBS. Live/dead cell labeling was performed using ZombieAqua (BioLegend, cat#423102), followed by staining with anti-puromycin antibody (12D10) for 20 minutes at 4°C. 2% rat serum (Stem cell) was used for Fc blocking. Stained cells were washed with 1 ml cold FACS buffer and resuspended in 200 μl. Surface receptor staining was analyzed using BD FACSCanto II.

#### Immunopeptidomics

##### Cells preparation

WT, STING^RRAA^, and STING^-/-^ RAW cells were seeded at a density of 100 million cells per plate one day prior to the experiment. Cells were either treated with LPS for 6 hours or left untreated. Following treatment, cells were scraped, pelleted by centrifugation at 1500 rpm for 5 minutes, flash-frozen in liquid nitrogen, and stored at -80°C for further processing.

##### Preparation of CNBR-activated Sepharose 4B beads

CNBR-activated Sepharose 4B beads (Sigma, cat#17-0430-01) were resuspended in 1 mM HCl at a ratio of 40 mg beads per 13.5 ml 1 mM HCl. The suspension was incubated with gentle tumbling at room temperature for 30 minutes. Beads were centrifuged at 215g for 1 minute at 4°C, and the supernatant was discarded.

##### Coupling of antibodies to beads

CNBR-activated Sepharose 4B beads (40 mg) were resuspended in 4 ml of coupling buffer (0.1 M NaHCO₃ / 0.5 M NaCl, pH 8.3), centrifuged at 215g for 1 minute at 4°C, and the supernatant discarded. Mouse antibodies Pan-H2 (BioXcell, cat#BE0077, RRID:AB_1125537), H2-Kb (BioXcell, cat#BE0172, RRID:AB_10949300), and H-2Kd/H-2Dd (BioXcell, cat#BE0180, RRID:AB_10950841) were coupled to the beads at a ratio of 1 mg antibody per 40 mg beads in coupling buffer. The reaction was incubated for 120 minutes with gentle tumbling at room temperature. After coupling, beads were centrifuged at 215g for 1 minute at 4°C, and the supernatant was discarded. Beads were blocked by resuspending in 1 ml of blocking buffer (0.2 M glycine) and incubating for 30 minutes with gentle tumbling at room temperature. Supernatants were discarded, and the beads were washed twice by centrifugation with PBS (pH 7.2). Finally, beads were resuspended to a final concentration of 1 mg antibody per ml PBS (pH 7.2) and stored at 4°C.

##### Immunoisolation of MHC 1-associated peptides

Frozen cell pellets (100 million cells per sample) were thawed and resuspended in 0.5 mL PBS (pH 7.2). Cells were solubilized by adding 1 mL of detergent buffer consisting of PBS (pH 7.2), 1% (w/v) CHAPS (Sigma, cat#C9426-5G), and Protease Inhibitor Cocktail (Sigma, cat#P8340-5mL). The suspension was incubated with gentle tumbling for 60 minutes at 4°C and then centrifuged at 16,600g for 20 minutes at 4°C. Supernatants were transferred to new tubes containing covalently cross-linked CNBR-Sepharose beads conjugated to Pan-H2 (0.5 mL), H2-Kb (0.333 mL), and H-2Kd/H-2Dd (0.333 mL) antibodies per sample. Samples were incubated with gentle tumbling for 180 minutes at 4°C. Beads were transferred to BioRad Poly-Prep chromatography columns, and the flowthrough was discarded. Beads were washed sequentially with PBS, 0.1X PBS, and water. MHC I complexes were eluted from the beads using 1% trifluoroacetic acid (TFA). Acidic filtrates containing peptides were separated from MHC I subunits, including HLA molecules and β-2 microglobulin, using homemade stage tips packed with two 1 mm diameter octadecyl (C-18) solid-phase extraction disks (EMPORE). Stage tips were pre-washed with methanol, followed by 80% acetonitrile (ACN) in 0.1% TFA, and then 1% TFA. Samples were loaded onto the stage tips, washed sequentially with 1% TFA and 0.1% TFA, and peptides were eluted with 30% ACN in 0.1% TFA. Eluates were dried using vacuum centrifugation and stored at -20°C until mass spectrometry analysis.

##### MS analysis

Samples were dried down and solubilized in 5% ACN-4% formic acid (FA). The prepared samples then loaded onto a homemade reversed-phase column (150-μm inner diameter, 200 mm length) with a 116-min gradient from 10 to 30% ACN-0.2% FA and a 600-nl/min flow rate on an Easy nLC-1200 (RRID:SCR_026978) coupled to a Exploris 480 (Thermo Fisher Scientific, RRID:SCR_027000). Each Full MS spectra were acquired at a resolution of 120,000, followed by data-dependent tandem-MS (MS/MS) scans of the most abundant multiply charged precursor ions within a 3-second acquisition window. MS/MS experiments utilized higher-energy collision dissociation (HCD) with a collision energy setting of 34%. The automatic gain control (AGC) target was set to 100%, with a resolution of 30,000 and an injection time of 1000 ms for the MS/MS scans.

##### Targeted LC-MS/MS analyses using parallel reaction monitoring (PRM)

For the targeted quantification of gB peptide in RAW cell extracts, immunoprecipitation MAPs were isolated from 100 million RAW cells for each condition. LC-MS/MS experiments were performed on the same instrument and elution conditions described above, except that MS survey scans were acquired at a resolution of 120,000, automatic gain control at 4 × 10^5^ and maximum injection time at 251 ms. Scheduled targeted HCD MS/MS scans were acquired at a resolution of 45,000 and used an isolation window of 1.2 *m*/*z* with 27% normalized collision energy. Skyline^69^ was used to extract the endogenous MS/MS spectrum of gB peptides.

##### Data processing

Database searches were performed using the PEAKS engine (Bioinformatics Solutions Inc., version XPro, https://www.bioinfor.com/peaks-xpro/, RRID:SCR_022841) against the mouse Uniprot database (20220406). Error tolerances for precursor mass and fragment ions were set to 10.0ppm and 0.01 Da, respectively. A non-specific digest mode was used, and variable modifications included phosphorylation (STY), Oxidation (M), and Deamidation (NQ). Peaks search results were subsequently processed using MAPDP https://capa.iric.ca/proteomics/resources ^70^, applying the following filters peptides of 8-15 amino acids in length, a rank-eluted ligand threshold ≤ 2% based on NetMHCpan-4.1b predictions, and a 5% false discovery rate (FDR). FDR was calculated using the decoy hits imported from Peaks, which employs a decoy-fusion strategy ^71^.

#### Transcriptomics

##### RNA extraction and RNA-Seq

Total RNA was isolated using the AllPrep DNA/RNA/miRNA Universal kit (QIAGEN, 7326820) following the manufacturer’s instructions. RNA concentration was determined using Qubit (Thermo Scientific), and RNA quality was assessed with the 2100 Bioanalyzer (Agilent Technologies). Transcriptome libraries were generated using the KAPA RNA HyperPrep (Roche, 7962363001) incorporating a poly-A selection (Thermo Scientific). Sequencing was performed on the Illumina NextSeq500, obtaining 15-20M single-end 84bp reads per sample. Library preparation and sequencing were conducted at the Genomics Platform of the Institute for Research in Immunology and Cancer (IRIC).

##### Mapping protocol: GRCm38 genome

Sequencing reads were trimmed for adapter sequences and low quality 3’ bases using Trimmomatic version 0.35 (http://www.usadellab.org/cms/index.php?page=trimmomatic, RRID:SCR_011848) ^72^. The cleaned reads were then aligned to the reference mouse genome version GRCm38 with gene annotation from Gencode version M24 (https://www.gencodegenes.org/mouse/release_M24.html, RRID:SCR_014966) (based on Ensembl 99) using STAR version 2.7.1a (https://www.encodeproject.org/software/star/, RRID:SCR_004463) ^73^.

##### Differential gene expression

Gene expressions were obtained both as raw read counts from STAR and as normalized gene and transcript-level expression in TPM values, computed using RSEM (RRID:SCR_000262) ^74^, for the stranded RNA libraries. Gene read counts were then normalized using DESeq2 version 1.30.1(https://bioconductor.org/packages/release/bioc/html/DESeq2.html, RRID:SCR_015687) ^75^. Differential gene expression analyses for the WT LPS vs WT no LPS comparison were also performed using DESeq2.

#### Quantification and Statistical Analysis

The results presented in this study are representative of at least three independent experiments. Statistical analyses were performed using GraphPad Prism 9, including Student’s t-test, one-way ANOVA with Dunnett’s post-test, and two-way ANOVA. A p-value of less than 0.05 was considered statistically significant and denoted by *, while p-values of less than 0.01, 0.001, and 0.0001 were marked as **, ***, and ****, respectively.

Differential analysis of immunopeptidome was performed using Limma Trend R package, http://bioinf.wehi.edu.au/limma/ (LIMMA, RRID:SCR_010943) ^76^, with default parameters. Intensity data, provided by PEAKS, were normalized using the variance stabilization normalization (vsn) method ^77^. Missing values were imputed using the MinProb method from imputeLCMD R package (https://cran.r-project.org/web/packages/imputeLCMD/index.html), with a quantile parameter (q) set to 0.05.

---

**Supp. Fig. 1:**
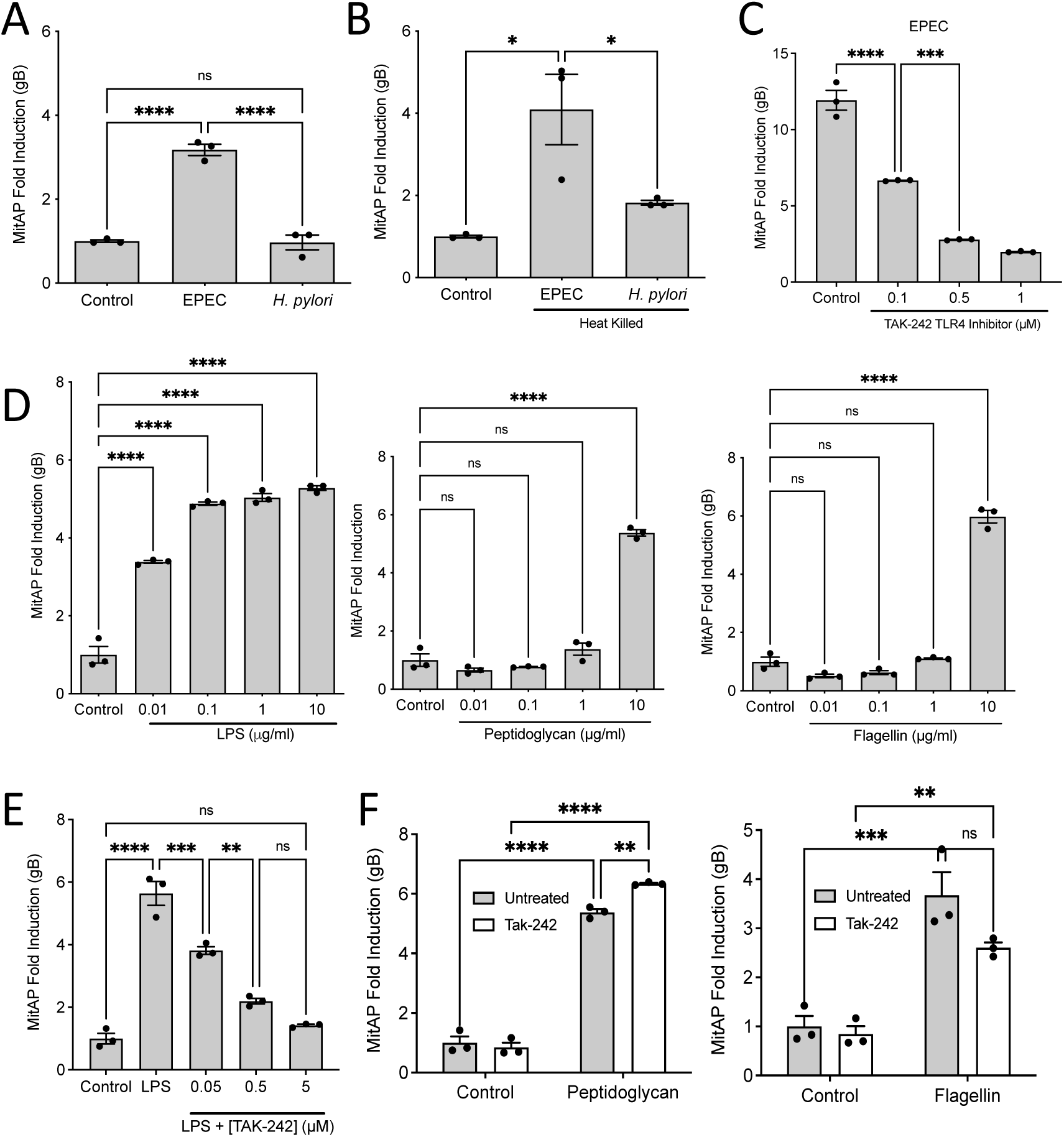
Gram-negative bacteria and LPS activate MitAP through TLR4. Related to Fig. 1. **A**) Of two Gram-negative bacteria, EPEC, but not *H. pylori*, activates MitAP. **B**) Similar results are observed with heat-killed bacteria, ruling out the need for an active inhibition. **C**) TAK242, a specific TLR4 inhibitor, inhibits the activation of MitAP by EPEC. **D**) Peptidoglycan and Flagellin activate MitAP only at a very high concentration, while LPS activates MitAP at a low concentration. **E**) TAK242 inhibits the activation of MitAP by LPS but fails to inhibit the activation of MitAP by peptidoglycan and flagellin (**F**).

**Supp. Fig. 2:**
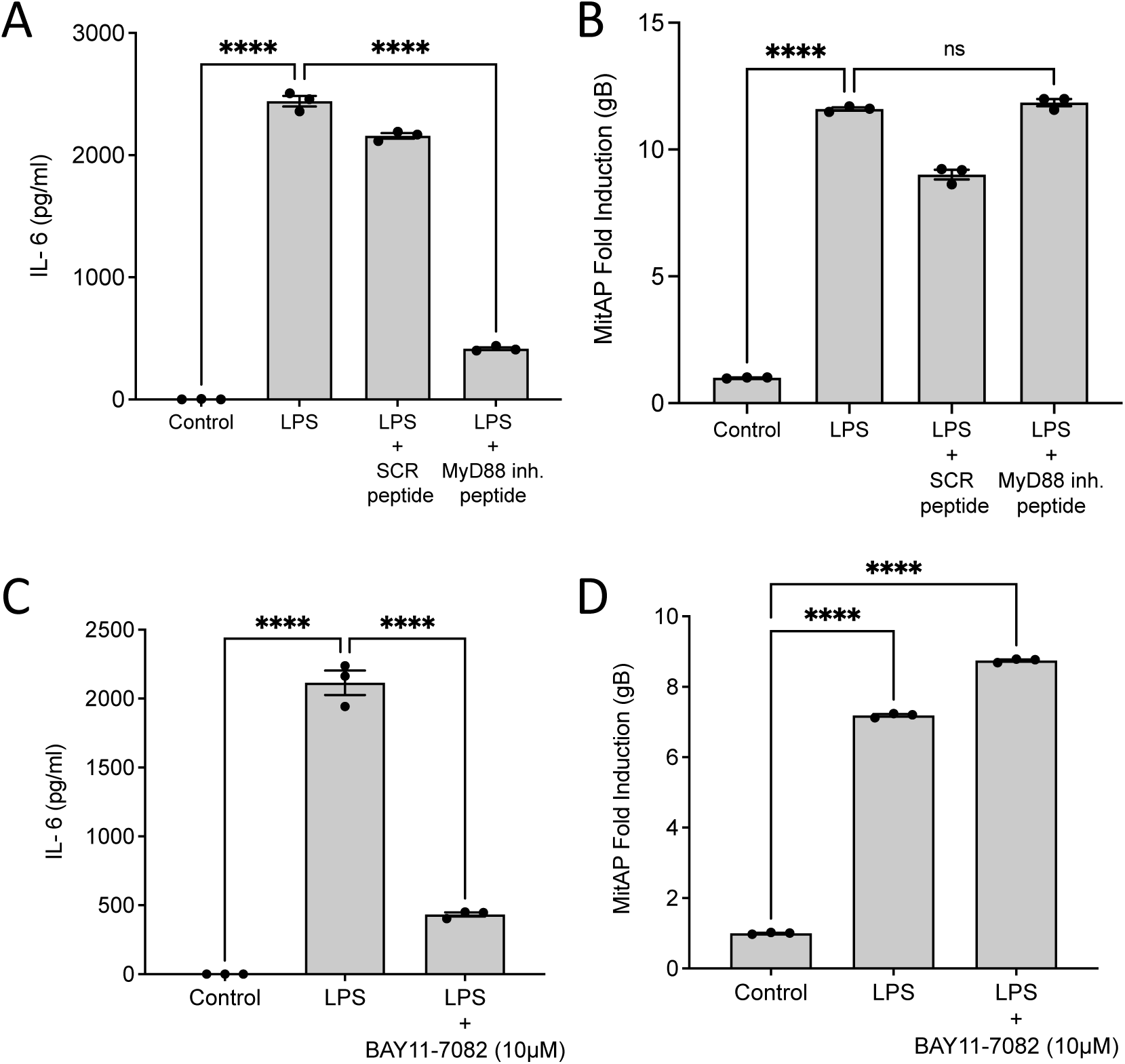
MitAP activation is MyD88-independent. Related to Fig. 1. **A**) Pre-treatment of RAW cells with a specific MyD88 inhibitory peptide inhibits the release of IL-6 in response to LPS. **B**) The activation of MitAP in response to LPS is not inhibited by the MyD88 inhibitory peptide. **C)** Pre-treatment of RAW cells with a specific NF-kB inhibitor (Bay11-7082) inhibits the release of IL-6 in response to LPS but not MitAP (**D**).

**Supp. Fig. 3:**
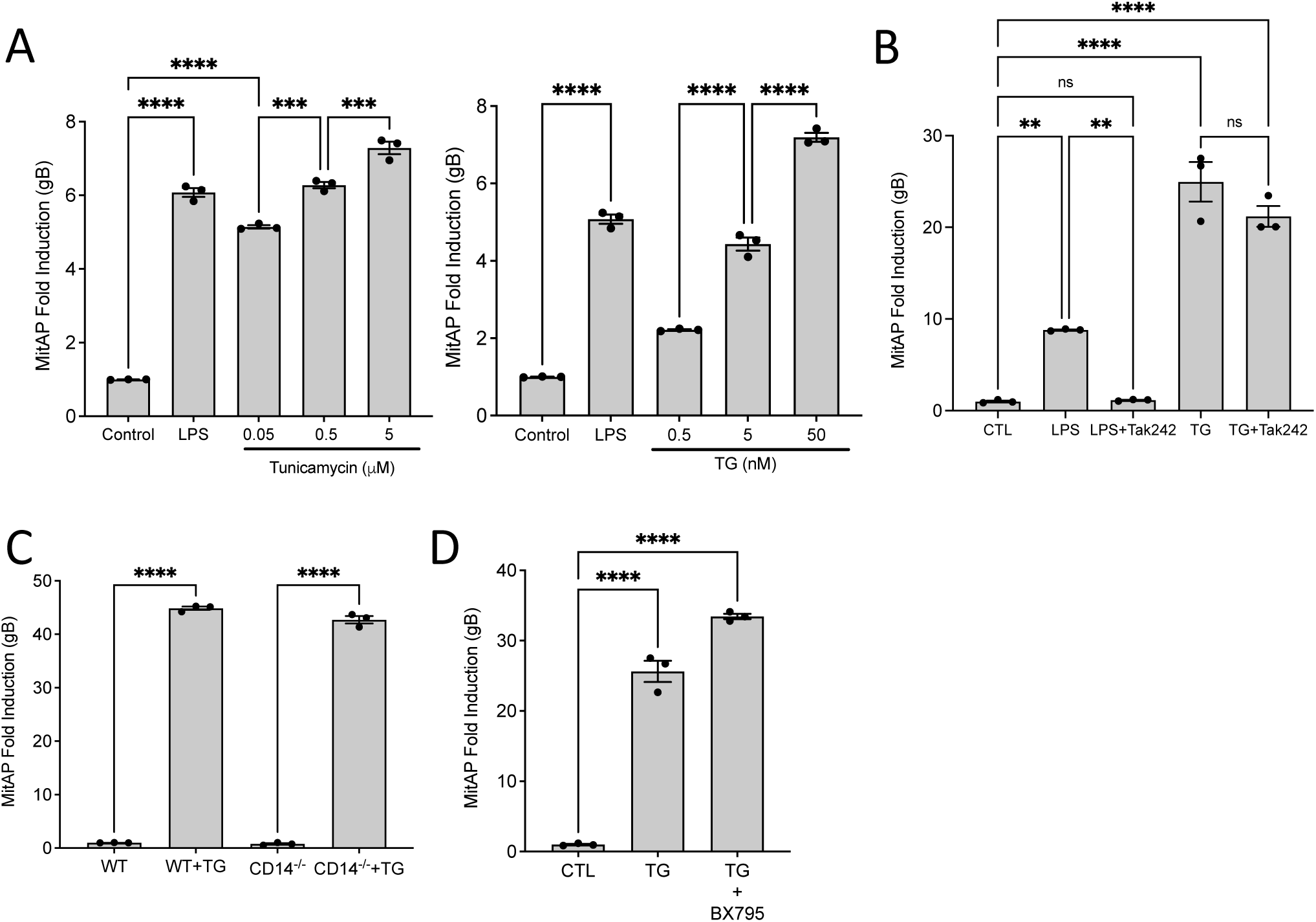
The UPR activates MitAP. Related to Fig. 4. **A**) Tunicamycin and Thapsigargin (TG) treatments induce MitAP in a dose-dependent manner in WT RAW macrophages. **B**) Activation of MitAP by TG is not inhibited by TAK242. **C**) Activation of MitAP by TG is not inhibited in the CD14^-/-^ RAW cells. **D**) Activation of MitAP by TG is not inhibited by BX795 (TBK1 inhibitor).

**Supp. Fig. 4:**
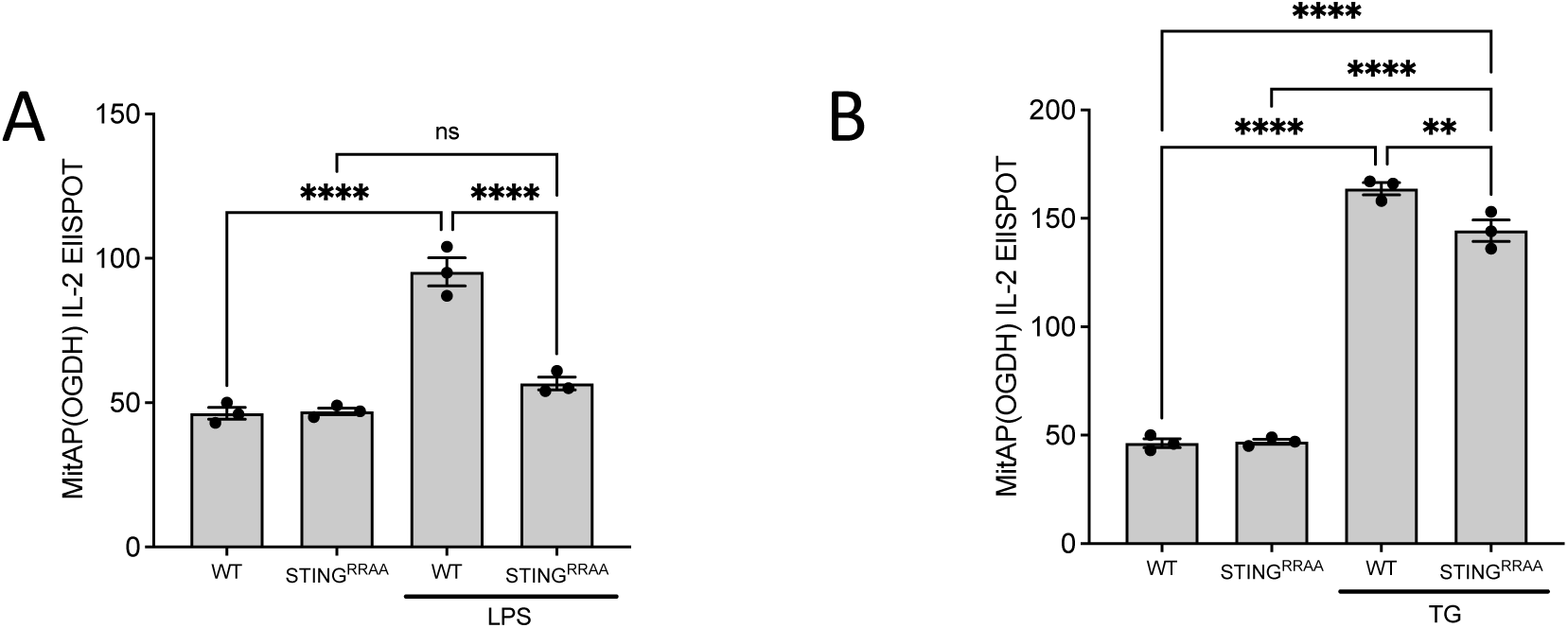
STING affects the presentation of a peptide from OGDH. Related to Fig. 4. **A**) The presentation of a peptide from the mitochondria matrix protein OGDH is inhibited in the STING^RRAA^ cells. **B**) Presentation of the OGDH peptide is efficiently activated by TG in STING^RRAA^ cells.

**Supp. Fig. 5:**
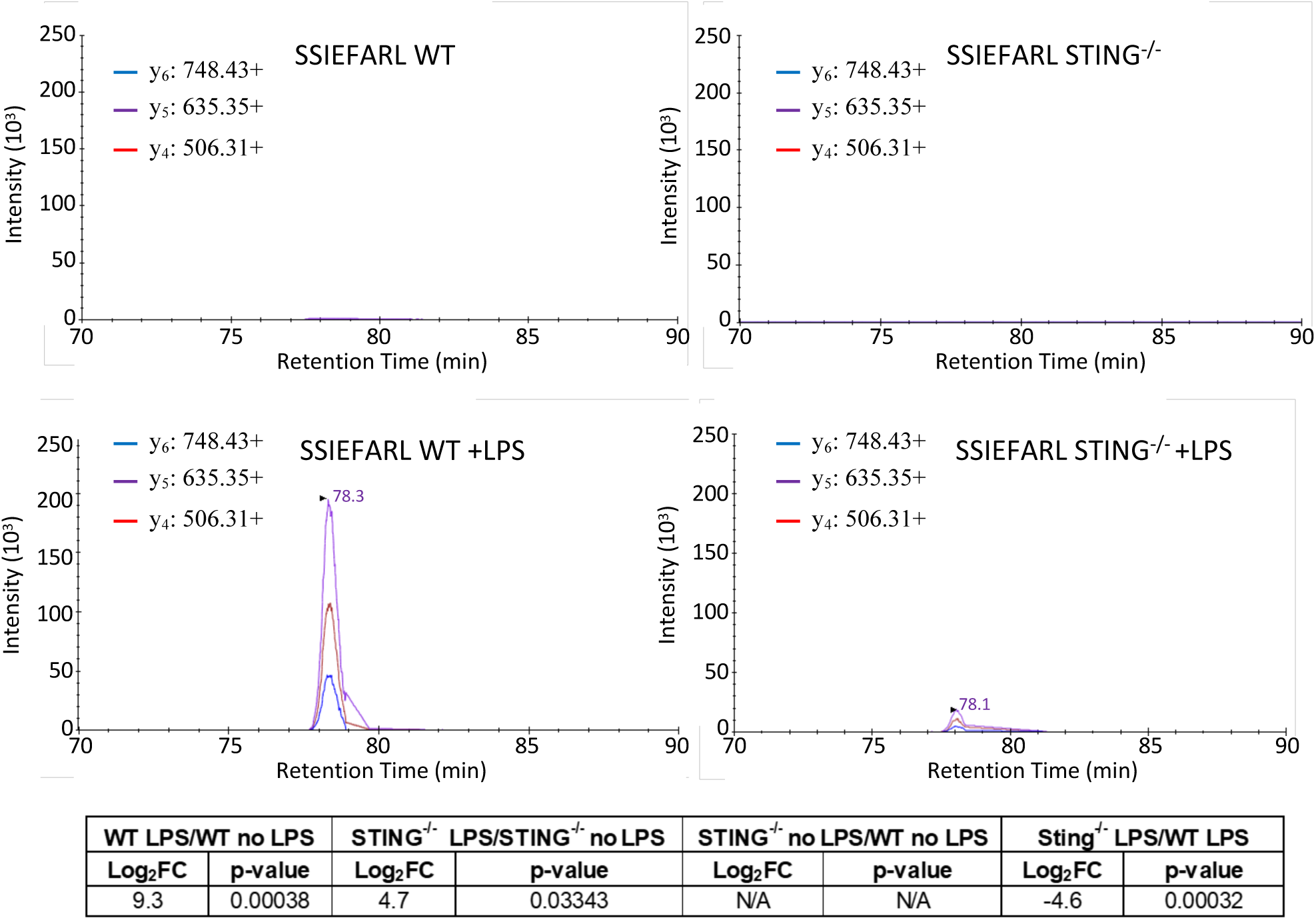
PRM confirms the STING-dependent induction of MitAP by LPS. Related to Fig. 4. Representative extracted ion chromatograms (XICs) of the SSIEFARL peptide (precursor m/z = 461.7533, z=2) using LC-MS/MS with parallel reaction monitoring (PRM) are shown for cells expressing WT or STING^-/-^ RAW macrophage cells, treated or not with LPS. The y-type fragment ions (y_4_: 506.31, y_5_: 635.35, y_6_: 748.43) were used for confirmation and quantification. The peptide was reproducibly detected at 78.5 min exclusively in LPS-treated WT cells, confirming its presentation via the MitAP pathway. The peptide was not detected in non-stimulated cells and was strongly reduced in STING^-/-^ cells despite LPS treatment. The value for the gB peptide induction is shown below.

**Supp. Fig. 6:**
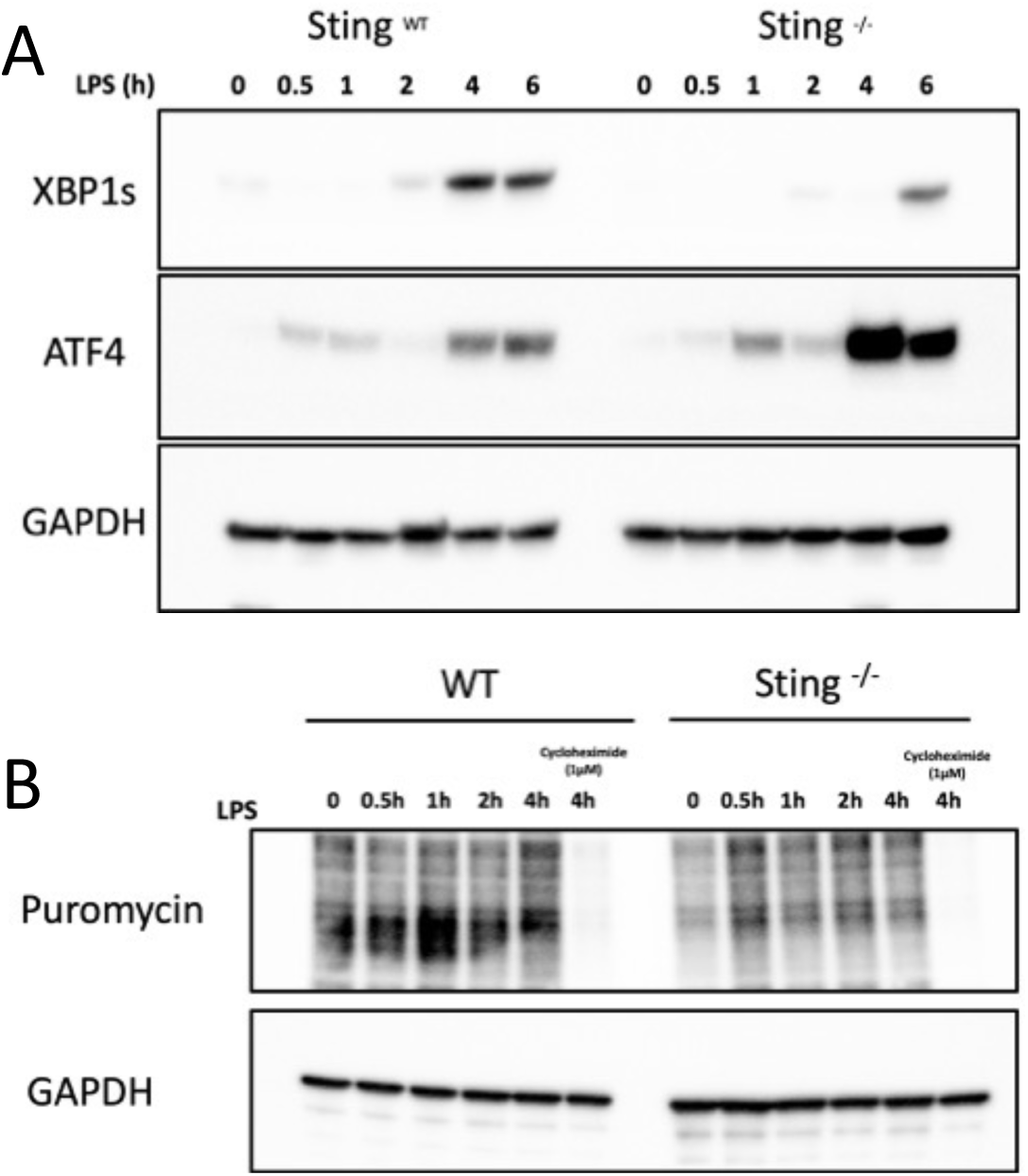
The loss of STING affects UPR activation in response to LPS. Related to Fig. 4. **A**) XBP1s and ATF4 are produced in response to LPS in WT RAW cells. The production of ATF4 is increased, and XBP1s is strongly decreased in the STING^-/-^ cells. **B**) Western blotting with anti-puromycin antibodies indicates that protein synthesis is strongly decreased in the STING^-/-^ cells. Protein synthesis is inhibited when cells are treated with Cycloheximide before LPS (negative control).

**Supp. Fig 7:**
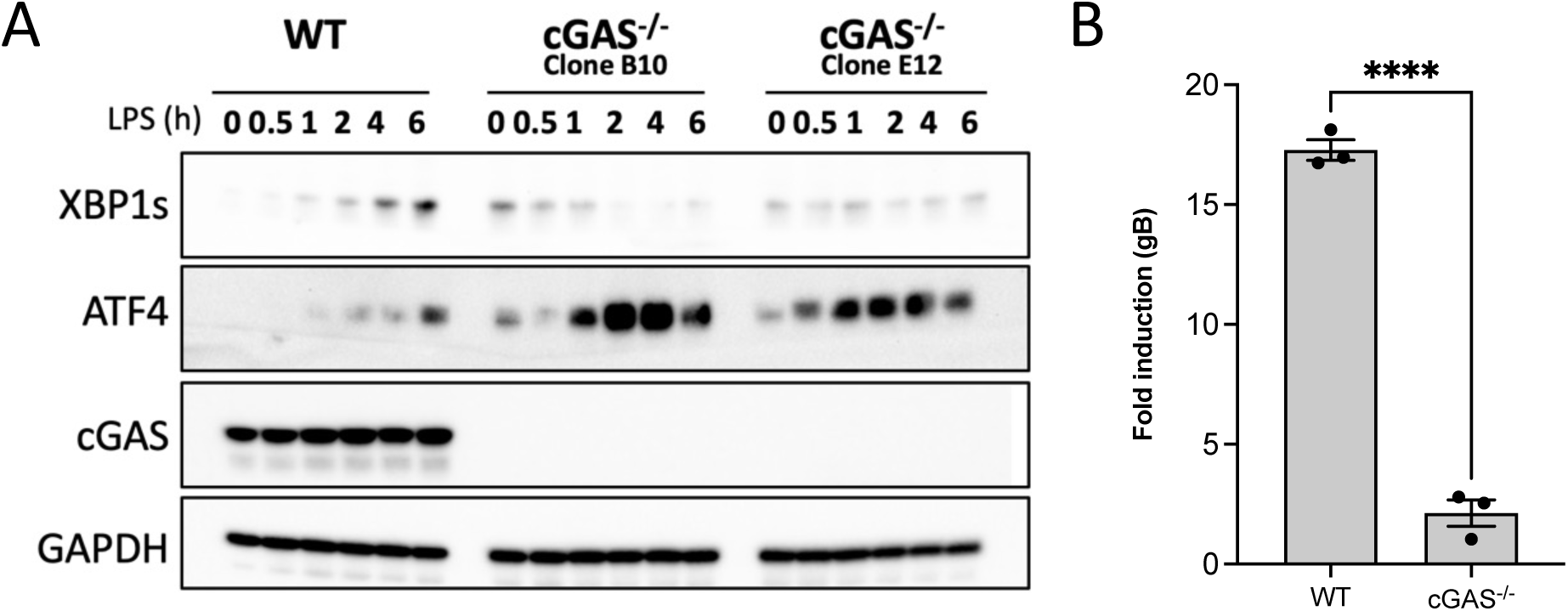
cGAS is required for MitAP and the UPR regulation. Related to Fig. 4. **A)** Both XBP1s and ATF4 are produced in response to LPS in WT RAW cells. In contrast, ATF4 is strongly increased, and the production of XBP1s is decreased in the cGAS^-/-^ cells. **B**) MitAP is inhibited in cGAS^-/-^ RAW cells in response to LPS.

**Supp. Fig 8:**
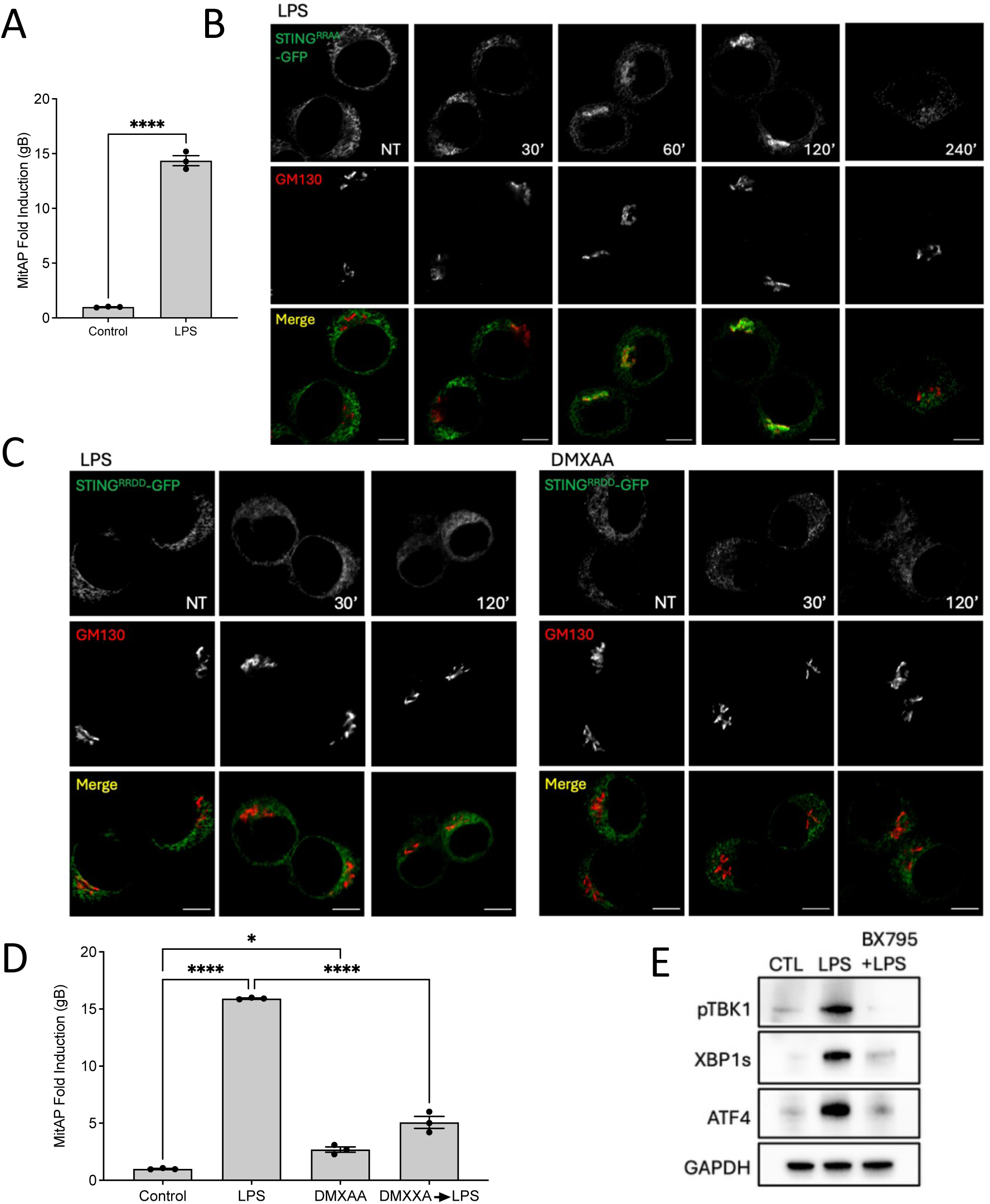
The “UPR motif” plays a role in the trafficking of STING out of the ER. Related to Fig. 4. **A**) LPS treatment induced a strong induction of MitAP. **(B)** LPS stimulation induces translocation of STING^RRAA^ to the Golgi, as assessed by anti-GFP staining and GM130 labeling, by 60 min, followed by exit from the Golgi by 240 min. Scale bar, 5 µm. **(C)** Neither LPS nor DMXAA treatment induces translocation of STING^RRDD^ to the Golgi up to 120 min. Scale bar, 5 µm. **D)** Pre-treatment of WT RAW cells with DMXAA, which induces STING translocation to the Golgi, attenuates LPS-induced MitAP. **E**) Pre-treatment of WT RAW cells with the TBK1 inhibitor (BX795) inhibits LPS-induced UPR.

**Supp. Fig. 9:**
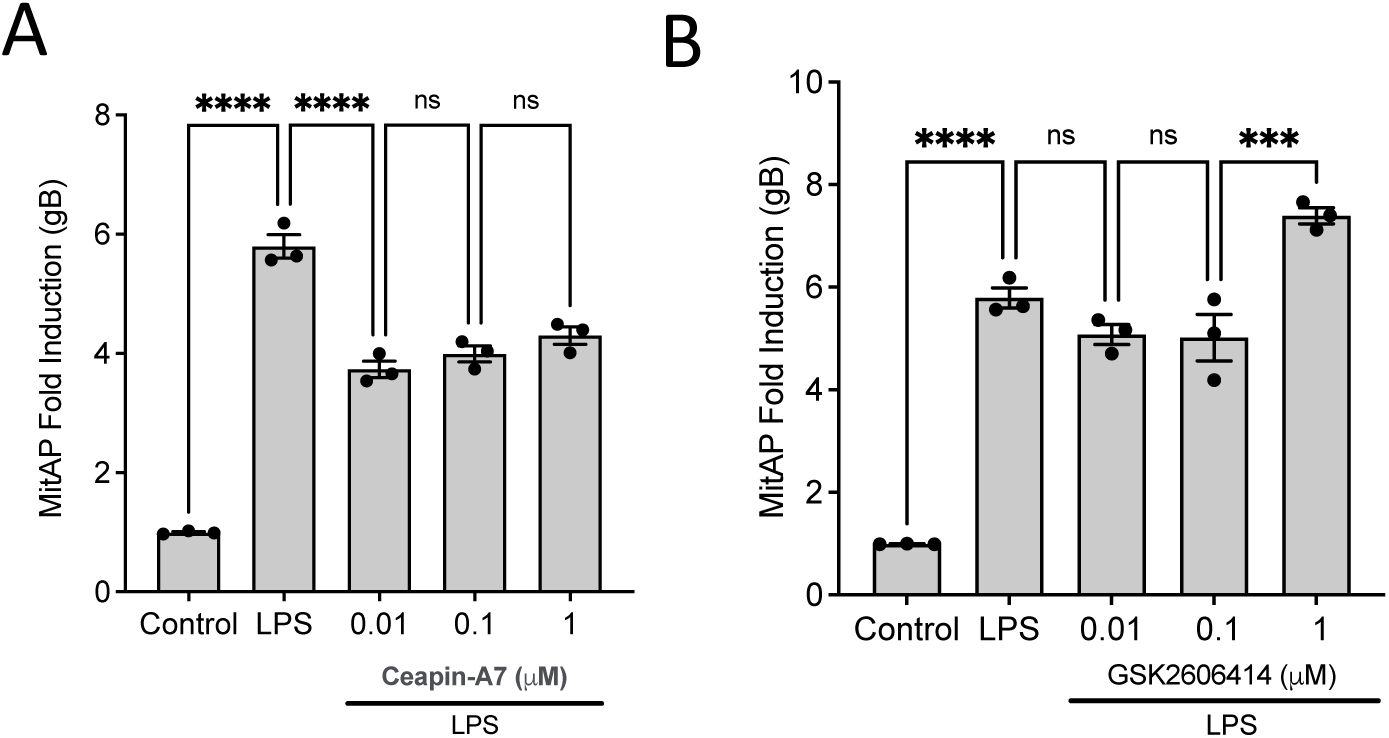
The inhibition of PERK and ATF6 does not affect MitAP activation by LPS. Related to Fig. 5. Treatment with Ceapin-A7 (ATF6 inhibitor) (**A**) and GSK2606414 (PERK inhibitor) (**B**) has no inhibitory effect on the level of MitAP induced by LPS.

**Supp. Fig. 10:**
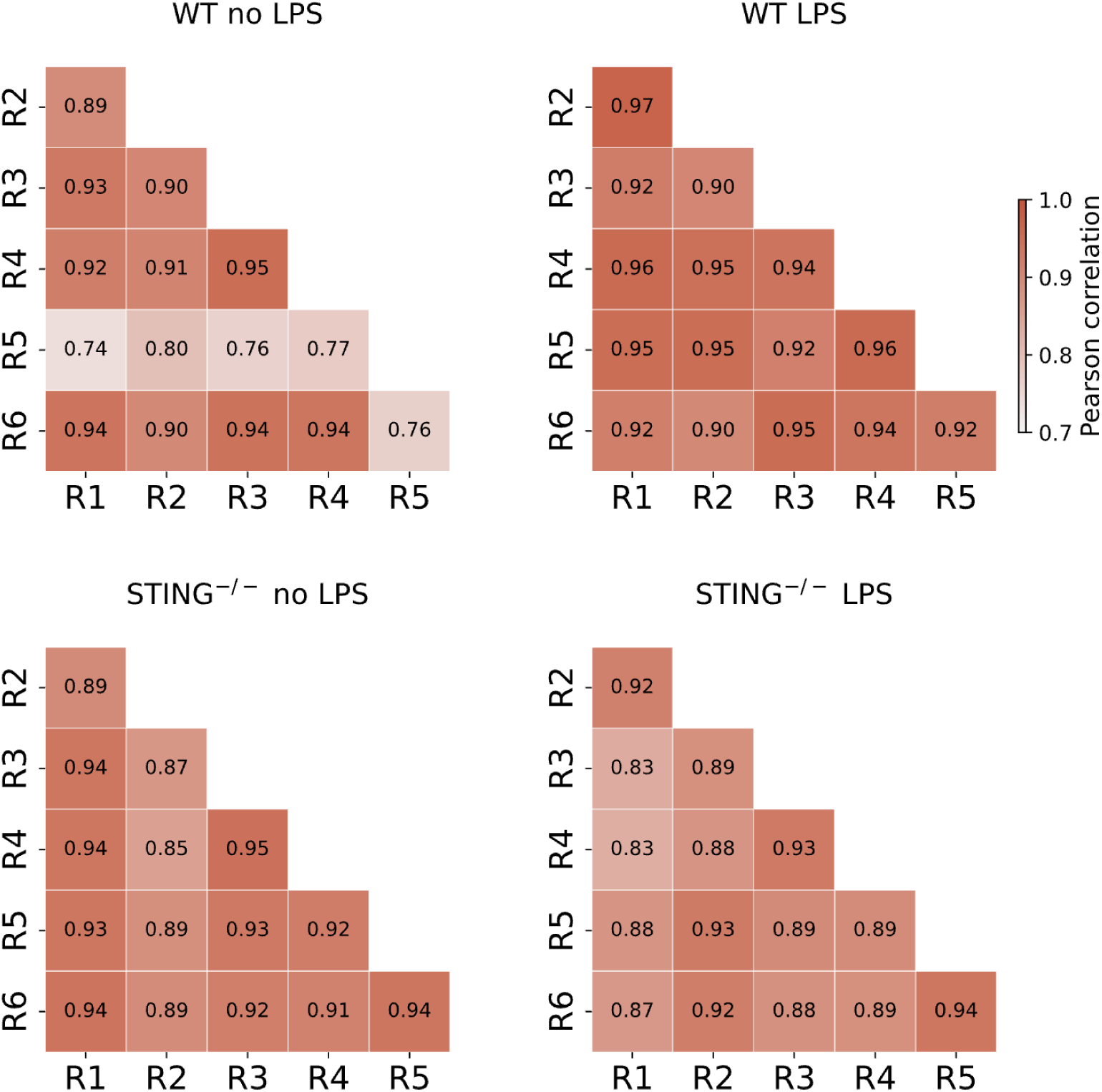
Peptides upregulated by LPS are processed from source proteins involved in immune functions. Related to Fig. 7. Gene Ontology (GO-terms) of the source proteins that had peptides presented in higher or lower amounts in response to LPS.

**Supp. Fig. 11:**
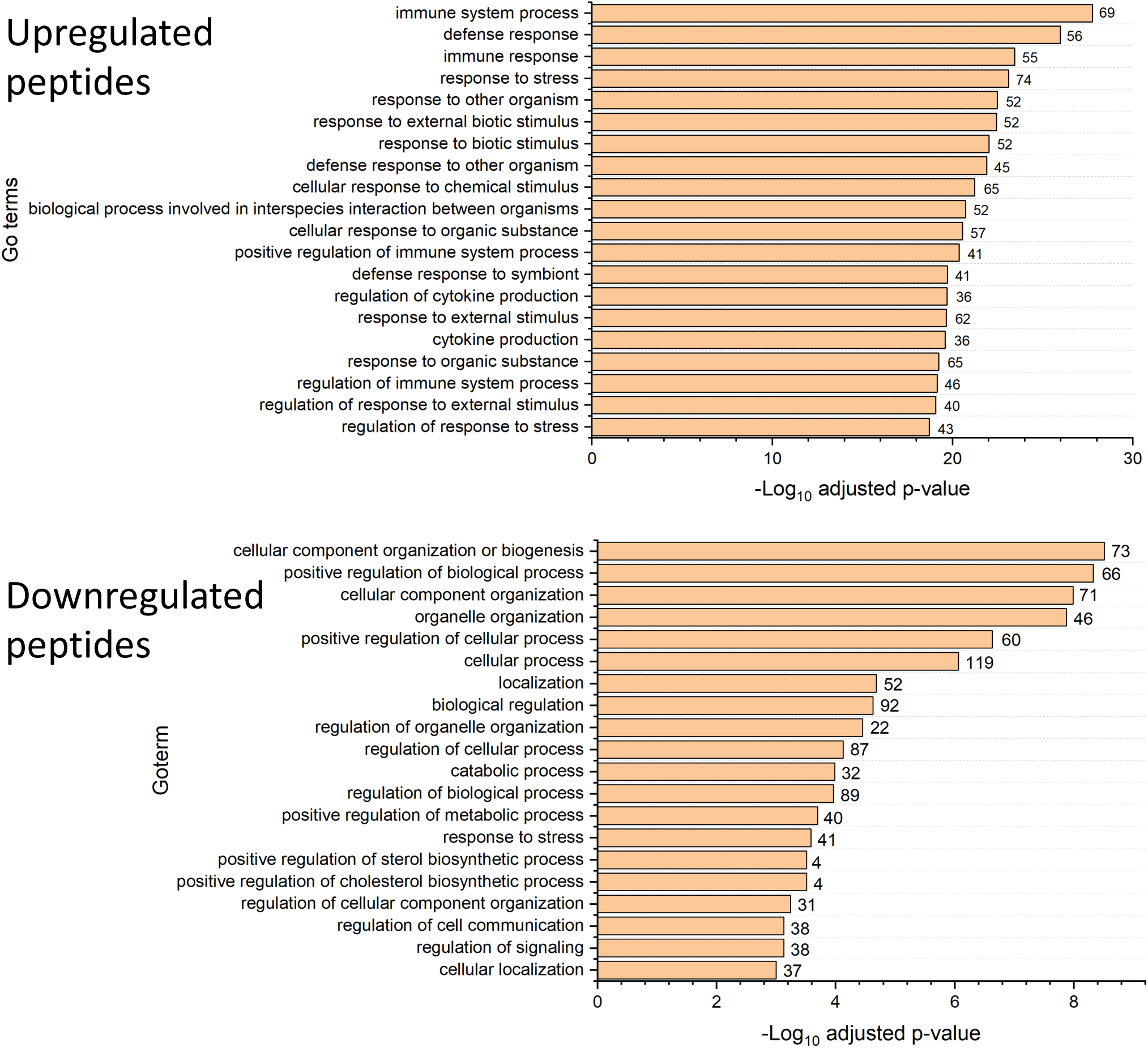
MHC-I peptide quantification is reproducible across replicates. Related to Fig. 7. Heatmaps display pairwise Pearson correlation coefficients of peptide intensities for six independent biological replicates (R1–R6) in each condition: WT no LPS (top left), WT +LPS (top right), STING⁻/⁻ no LPS (bottom left), and STING⁻/⁻ +LPS (bottom right). Correlations were computed using normalized peptide intensities across all detected MAPs. Strong intra-group correlations in the WT + LPS condition (r > 0.92) demonstrate high reproducibility of LPS-induced antigen presentation.

**Supp. Fig. 12:**
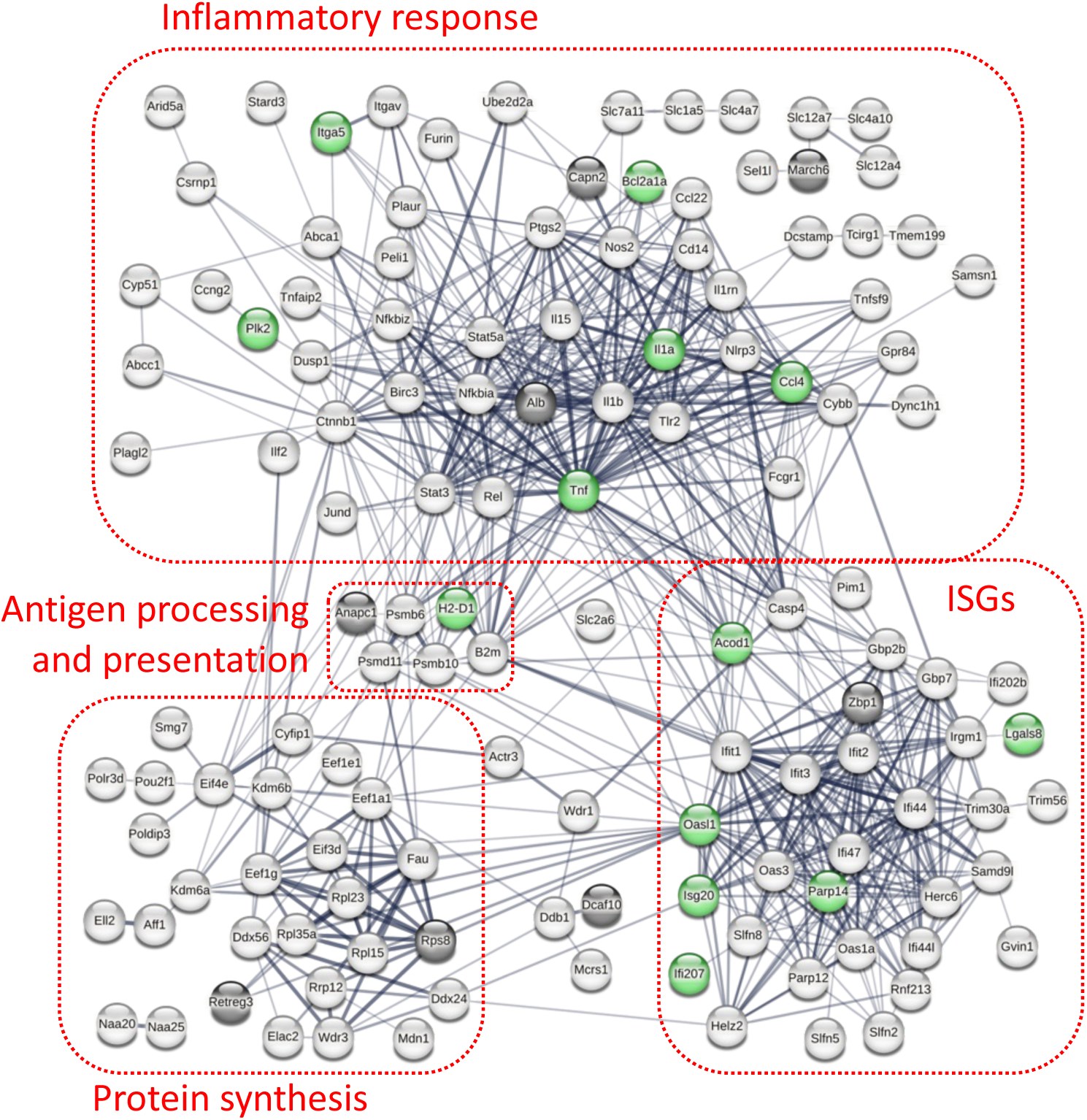
Effect of the loss of function of STING “UPR motif” on self-peptide presentation. Related to Fig. 7. STRING interactome of the source proteins that had peptides presented in higher amounts (> 5-fold) in response to LPS (WT +LPS/WT untreated). Proteins in black are the source of peptides whose presentation is down-regulated by at least 2-fold in response to LPS in STING^RRAA^ cells (STING^RRAA^ +LPS/WT +LPS). Proteins in green are upregulated by at least 2-fold. The edges indicate functional and physical protein associations. The line thickness indicates the strength of the data support. The assignment in the red boxes was based on a literature survey.

Supp. Table 1: **List of peptides identified by mass spectrometry.** An Excel file shows the peptide data.

**Supp. Table 2:** The STING-UPR axis regulates the presentation of a set of antigens. Related to Fig. 7. The presentation of the gB and OGDH peptides associated with the MitAP pathway (mitochondria) is inhibited in both the STING^-/-^ and STING^RRAA^ cells. A restricted set of peptides originating from protein sources present in other organelles is also presented by this pathway.

| Peptide sequence | Gene name | Mutant LPS vs WT LPS |  |  |  |
| --- | --- | --- | --- | --- | --- |
|  |  | Sting<sup>-/-</sup> |  | Sting<sup>RRAA</sup> |  |
|  |  | log<sub>2</sub>FC | p.value | log<sub>2</sub>FC | p.value |
| GYLLILHEL | Dcaf10 | -1.7 | 0.033 | -3.0 | 0.026 |
| RTYIFTFL | Rasgef1a | -1.0 | 0.006 | -4.6 | 0.014 |
| SPVRAELL | Mcmbp | -1.4 | 5.87223E-05 | -1.6 | 0.046 |
| SSLKFLLL | Ddx11 | -2.4 | 0.016 | -2.1 | 0.045 |
| TSLRFVFL | Retreg3 | -1.5 | 0.026 | -3.4 | 0.020 |
| VGAERNVLI | Hspa8 | -1.8 | 0.013 | -2.4 | 0.016 |
| VTLVFEHI | Cdk4 | -1.4 | 0.015 | -2.4 | 0.039 |
| VYAPLKELL | Ndc80 | -1.0 | 0.001 | -2.9 | 0.043 |
| YVPSKSKI | Sbno2 | -0.8 | 0.018 | -2.0 | 0.049 |

